# Evolutionary responses of a reef-building coral to climate change at the end of the last glacial maximum

**DOI:** 10.1101/2022.04.02.486852

**Authors:** Jia Zhang, Zoe T. Richards, Arne A. S. Adam, Cheong Xin Chan, Chuya Shinzato, James Gilmour, Luke Thomas, Jan M. Strugnell, David J. Miller, Ira Cooke

## Abstract

Climate change threatens the survival of coral reefs on a global scale, primarily through mass bleaching and mortality as a result of marine heatwaves. While these short-term effects are clear, predicting the fate of coral reefs over the coming century is a major challenge. One way to understand the longer-term effects of rapid climate change is to examine the response of coral populations to past climate shifts. Coastal and shallow-water marine ecosystems such as coral reefs have been reshaped many times by sea-level changes during the Pleistocene, yet, few studies have directly linked this with its consequences on population demographics, dispersal, and adaptation. Here we use powerful analytical techniques, afforded by haplotype phased whole-genomes, to establish such links for the reef-building coral, *Acropora digitifera*. We show that three genetically distinct populations are present in northwestern Australia, and that their rapid divergence since the last glacial maximum (LGM) can be explained by a combination of founder-effects and restricted gene flow. Signatures of selective sweeps, too strong to be explained by demographic history, are present in all three populations and overlap with genes that show different patterns of functional enrichment between inshore and offshore habitats. In contrast to rapid divergence in the host, we find that photosymbiont communities are largely undifferentiated between corals from all three locations, spanning almost 1000 km, indicating that selection on host genes and not acquisition of novel symbionts, has been the primary driver of adaptation for this species in northwestern Australia.

## Introduction

Glacial cycling during the Pleistocene is thought to be a major driver of biodiversity dynamics (Hewitt 2000; Hofreiter and Stewart 2009) and its effects provide important lessons that can be used to help predict the impacts of future climate change (Hofreiter and Stewart 2009; Nogués-Bravo et al. 2018). Population genetics is a valuable tool to understand these past climate events because it can reveal historical changes in species’ demography, connectivity, and diversity. Widespread application of population genetic tools to terrestrial (Hofreiter and Stewart 2009) and marine (Mattingsdal et al. 2019) species in the northern hemisphere has revealed a predominant picture of persistence in southern refugia followed by expansion and northward migration after the last glacial maximum (LGM), with more recent work describing differential species’ responses depending on habitat requirements (Hofreiter and Stewart 2009) and patterns of dispersal (Mattingsdal et al. 2019). Much less is known about the impacts of past climate shifts on tropical marine systems such as coral reefs, despite the profound impacts that changes in temperature and sea-level would have had on these shallow-water marine habitats (Wilson 2013; Ludt and Rocha 2015; Webster et al. 2018).

Throughout the tropics, the dominant effect of low sea levels during the last glacial maximum was a dramatic reduction in the amount of shallow water habitat (Kleypas 1997; Ludt et al. 2015). In broad agreement with this, many studies across a range of coral reef taxa have observed signatures of recent population expansion (Crandall et al. 2008; Crandall et al. 2012; Delrieu-Trottin et al. 2017), however not all populations follow this pattern. Genome-wide approaches are now revealing differential demographic histories of cryptic and recently diverged populations (Bierne et al. 2003; Cooke et al. 2020; Underwood et al. 2020; Bongaerts et al. 2021), some of which show signatures of recent isolation and decline (Moran et al. 2019). Moreover, the ranges of diverged populations in the marine environment are sometimes difficult to reconcile with modern geography and potential for physical dispersal (Bierne et al. 2003; Cooke et al. 2020; Underwood et al. 2020; Bongaerts et al. 2021), and they may be better understood with reference to historical connectivity such as during past glacial maxima. A historical perspective may therefore be crucial to understanding gene flow and adaptation in extant populations. However, the value of this approach depends heavily on the temporal resolution of demographic analyses so that their timing can be linked to specific climate events, and with the ability to detect and characterise signatures of selection so that these can be used to assess modes of local adaptation.

Emerging techniques based on the sequentially Markovian coalescent (SMC) can be used to reconstruct demographic histories of species in unprecedented detail, potentially revealing links with past climate (Nadachowska-Brzyska et al. 2015; Kozma et al. 2016; Chattopadhyay et al. 2019; Lucena-Perez et al. 2020). However, the most widely used variant of this technique, PSMC (Li and Durbin 2011), has limited power to infer recent events, a problem exacerbated by large effective population size (Schiffels and Durbin 2014). Since corals and many other broadcast-spawning marine taxa have large effective population sizes, most studies so far have focussed on changes in the distant past that cover many glacial cycles (Prada et al. 2016; Mao et al. 2018; Fuller et al. 2020). Inferences within the timeframe of the most recent glacial cycle require more sophisticated methods such as MSMC (Schiffels and Durbin 2014) and SMC++ (Terhorst et al. 2016) that make use of larger datasets (multiple whole genomes) to improve the sampling of haplotypes that share a recent common ancestor.

Even in systems where the effects of past climate change on biodiversity are relatively well understood, the role of natural selection and adaptation in response to climate change remains uncertain (Nogués-Bravo et al. 2018). Addressing this gap for climate-sensitive taxa such as corals is a pressing issue (Torda et al. 2017) directly relevant to their conservation and management in the Anthropocene. Adaptive evolution in corals is complex because it is likely to involve selection on the coral hosts themselves, as well as selection on and/or exchange of their dinoflagellate photosymbionts. Symbiont exchange is of particular interest because it may enable corals to adapt rapidly to anthropogenic climate change (Berkelmans and Oppen 2006; Torda et al. 2017). Numerous studies have observed variation in host-symbiont associations along environmental gradients (Bongaerts et al. 2013; Camp et al. 2020; Ros et al. 2021), and experiments have demonstrated that a switch in symbiont partnership can be induced by stress (Matsuda et al. 2022). Another potential mode of climate adaptation in corals is selection on the coral host. A range of studies examining population genetic, and gene expression differences between heat-adapted and naive corals all suggest that adaptation to heat is likely to involve many loci (Palumbi et al. 2014; Dixon et al. 2015; Fuller et al. 2020; Thomas et al. 2021). Modelling efforts have also attempted to describe the envelope of population genetic parameters, and rate of climate change under which corals could adapt based on natural selection (Matz et al. 2018). So far, however, there are few studies (except see Smith et al. 2022) that identify signatures of selection in relation to adaptation and survival over a sustained period of warming, such as the transition from the LGM to today.

In this study, we used a population whole-genome sequencing approach to understand the impacts of past climate change on the widespread reef building coral, *A. digitifera* in northwestern Australia. In this region, *A. digitifera* is common on offshore atolls at the shelf-edge and also forms part of a diverse inshore community (in the Kimberley region) that thrives despite extreme heat, frequent aerial exposure and highly variable turbidity (Richards et al. 2015; Richards et al. 2019). Modern coral reefs in the Kimberley were extirpated during the last glacial maximum (LGM), while those offshore may have persisted but would have experienced a period of much reduced shallow-water habitat and been much closer to the coast (Wilson 2013; Solihuddin, O’Leary, et al. 2016; McCaffrey et al. 2020). The contrasting biogeography of these sites provides an ideal case study of the effects of climate change during the last glacial cycle, and our analytical approach is designed to investigate this comprehensively. We do so through demographic modelling based on multiple whole genomes providing accurate inferences in the window leading up to and following the LGM (1kya - 100kya), and through sensitive detection of signatures of recent selection via extended haplotype homozygosity and population branch statistics. In addition, we use non-host reads to profile the dinoflagellate symbionts inhabiting each coral colony based on standard markers such as the ITS2 region of ribosomal RNA as well as via mitochondrial sequences and a novel *k*-mer-based distance metric. This combination of approaches allows us to examine the interplay between demographic change, connectivity, selection and shifts in symbiont community composition during a rapid climate change event for the first time in a coral.

## Results

Whole-genome sequencing of 75 *Acropora digitifera* colonies from three reef systems in northwestern Australia yielded a mean per-sample coverage of 19.5X that we used to call approximately 9.6 million high-quality biallelic single nucleotide polymorphisms (SNPs) with GATK (supplementary fig. S1, supplementary table S2). Of the few coral whole-genome studies conducted to date, most (Shinzato et al. 2015; Cooke et al. 2020; Thomas et al. 2021) adopted a shallow sequencing approach (except see Fuller et al. 2020). The relatively high sequencing depth in our study allowed us to reliably call genotypes at more than 95% of sites in 90% of samples (supplementary fig. S1) supporting population-based haplotype phasing with SHAPEIT (Delaneau et al. 2012). As SHAPEIT infers missing genotypes based on phasing information, we tested its accuracy by removing genotypes with high-quality calls and then comparing their original value with that imputed by SHAPEIT. This confirmed that imputation (and by extension phasing) was generally highly accurate, relatively unaffected by minor allele frequency, but slightly better for sites with fewer missing values and for homozygous genotypes (supplementary fig. S2).

### Population structure in the coral host

PCA, and fineSTRUCTURE analysis (fig. 1C, supplementary fig. S3), showed clear genetic structure that divided corals from the six sampled reefs into three geographically separated groups, hereafter called North Offshore (NO) which includes Ashmore Reef, South Offshore (SO) which includes all reefs from the Rowley Shoals, and Inshore (IN) which includes two locations within macrotidal coral communities in the Kimberley (Adele Island, Beagle Reef). Using fineSTRUCTURE we also identified substructure within the inshore population between samples from Adele Island (AI) and Beagle Reef (BR) (supplementary fig. S3), however, the very tight clustering of all inshore samples in PCA analyses (PCs 1-3) indicated that this comprised a relatively minor component of genetic variation, and we therefore focussed on the three major clusters for our remaining analyses. Pairwise relatedness estimates based on shared genomic regions that were identical by descent (IBD) clearly partitioned samples into the three major clusters but failed to identify a distinction between BR and AI locations (supplementary fig. S4).

**Fig. 1.**
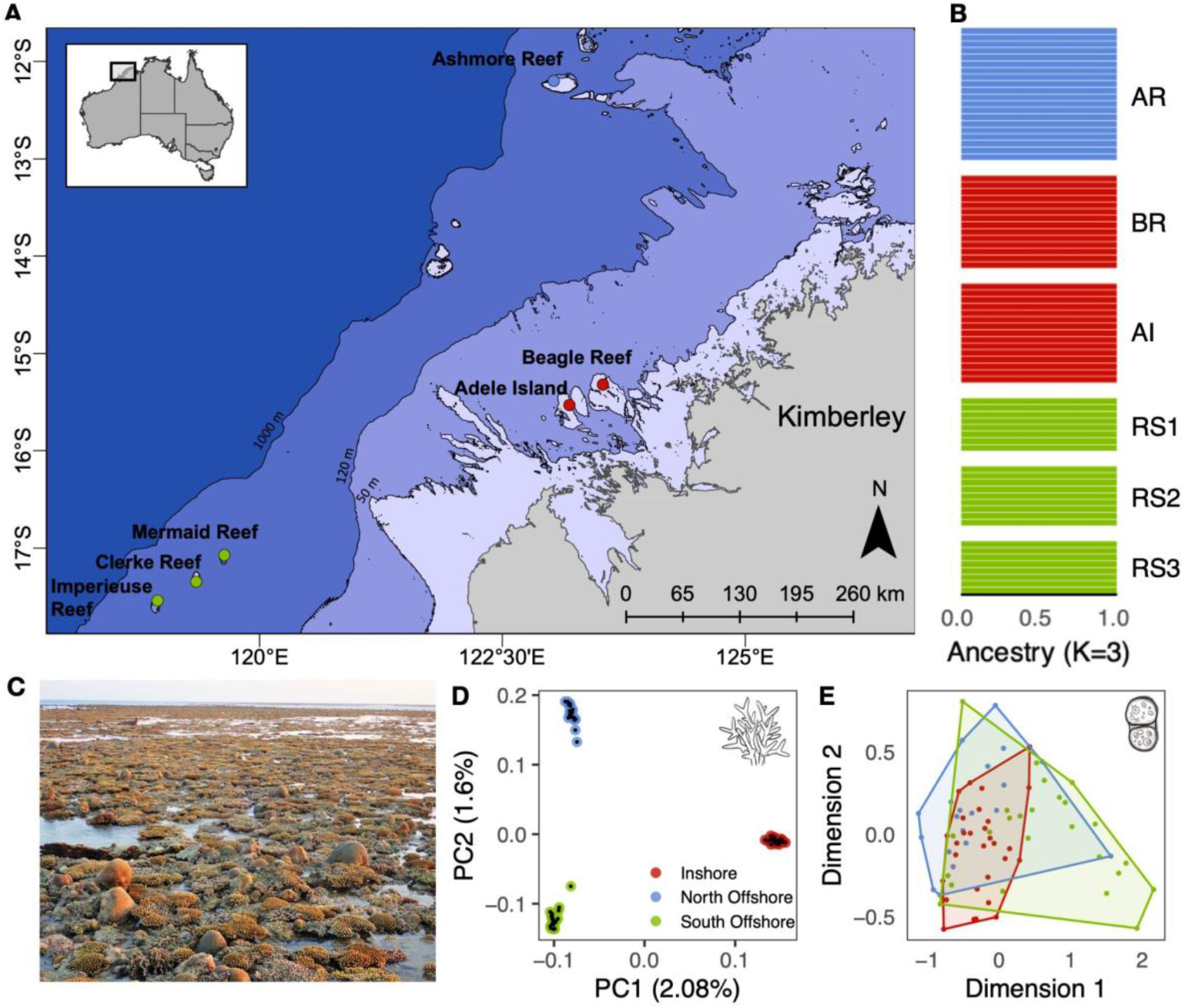
Sampling locations and genetic structure for the coral host and symbionts. All plots use the same colour scheme for locations as follows; North offshore, Ashmore Reef (AR) is shown in blue, inshore locations, Adele Island (AI) and Beagle Reef (BR) are shown in red, south offshore locations, Rowley Shoals (RS1: Mermaid Reef, RS2: Clerke Reef, RS3: Imperieuse Reef) are shown in green. **A**. Sampling locations in the Kimberley region, northwestern Australia. Bathymetric contours are shown at 50, 120, and 1000m depth with the present day landmass shown in grey. **B**. Admixture proportions for each colony calculated using ADMIXTURE with K=3 and coloured by the dominant cluster in each location. Each horizontal bar represents a single coral colony. **C**. Photograph of the reef flat at Adele Island showing corals exposed at low tide. Subaerial exposure for up to three hours during spring low tide is a characteristic feature of the inshore locations, AI and BR in this study. **D**. PCA showing the first and second principal components of genetic variation in the coral host. Points represent individual samples and are coloured by location. **E**. Multidimensional scaling plot showing relative pairwise distances between samples based on shared *k*-mers (d2s metric) from reads mapping to the dominant symbiont genus, *Cladocopium*. Convex hulls enclose points representing samples from the same location.

The relative distance between PCA clusters, a tree inferred by fineSTRUCTURE (supplementary fig. S3), and relative amounts of IBD segments indicated a closer relationship between the two offshore populations than between offshore and inshore. Consistent with this, genome-wide estimates of Fst were markedly lower (Fst ~ 0.007) between offshore populations than between north-offshore and inshore (Fst ~ 0.02) and south-offshore and inshore (Fst ~ 0.02). Despite low overall divergence (as measured with genome-wide Fst) between these populations, admixture coefficients (calculated using ADMIXTURE; Alexander et al. 2009) showed complete assignment (>99%) of each individual to its parent cluster (fig. 1B), suggesting that migration is rare or non-existent between locations. Demographic modelling with fastsimcoal2 (see below) confirmed this as it supported a model with recent gene flow but with very low migration coefficients (probability of migration/individual/generation ~1e-4; supplementary table S8). Analysis of simulated data under this model with ADMIXTURE produced the same complete assignment to locations as observed for the real data (supplementary fig. S15).

To place these Western Australian populations in a broader context we downloaded publicly available data from 16 *A. digitifera* colonies sampled from Okinawa, Japan (NCBI Bioproject PRJDB4188; Shinzato et al. 2015) and used PCAngsd (Meisner and Albrechtsen 2018), which is insensitive to differences in sequencing depth, to analyse them together with our samples. This confirmed that Japanese *A. digitifera* form a fourth genetic cluster, distinct from the three populations identified here (supplementary fig. S5) but with a level of divergence to offshore Western Australian populations that was similar to that between inshore and offshore clusters within Western Australia (Weighted Fst calculated with ANGSD, supplementary table S3B; cluster distance in PCA; supplementary fig. S5). In support of this low divergence, we also found that all four populations shared a single dominant mitochondrial haplotype (supplementary fig. S6) with few samples showing any variation from it. Finally, a phylogenetic tree based on established markers for phylogenetic inference in *Acropora* (Cowman et al. 2020) *confirmed that all four populations are likely conspecifics and congruent with the published A. digitifera* genome.

### Symbiont profiles

Based on the relative proportion of reads classified as Symbiodiniaceae by Kraken (Wood and Salzberg 2014) all samples from all locations were dominated by symbionts from the genus *Cladocopium* (supplementary fig. S8) which is the most common and diverse genus of symbiont in Indo-Pacific corals (LaJeunesse et al. 2018). To investigate the symbiont diversity within *Cladocopium*, we used three complementary approaches, all of which indicated that there was little difference in symbiont composition between locations. Firstly, a haplotype network based on consensus mitochondrial sequences (supplementary fig. S9B) for 41 samples where there was sufficient data (at least 20X mapping depth at mappable sites) revealed that all but one of the 41 samples were dominated by a single haplotype. This represents a much lower level of diversity than was observed in a previous study using the same approach to profile symbionts in *A. tenuis* on the GBR (Cooke et al. 2020). Since mitochondrial genomes are rarely used to profile Symbiodiniaceae (Waller and Jackson 2009; Gagat et al. 2017), and cannot easily be linked to known types, we also mapped the putative symbiont reads to the more-commonly used phylogenetic marker of ITS2 sequences, using the SymPortal database (Hume et al. 2019). This revealed a single ITS2 type profile comprising C40c, C72, C40, and C40e which occurred in most coral samples (supplementary fig. S9A). Finally, in order to minimise inherent biases in ITS2 or mitochondrial markers, we adopted an alignment-free approach based on analysis of shared *k*-mers (i.e. short sub-sequences of defined length *k*) (Reinert et al. 2009; Chan et al. 2014) in the symbiont reads to calculate a distance measure between all possible pairs of samples (see methods). An MDS plot based on this metric (fig. 1D) revealed similar levels of within-location to between-location diversity, confirming that there were no consistent differences in symbiont composition between locations.

### Demographic history and divergence times

To explore changes in effective population size (Ne) and to estimate divergence times among the coral populations identified above, we performed demographic modelling using two complementary approaches, SMC++ (Terhorst et al. 2016) and fastsimcoal2 (Excofffier et al. 2021). Translating demographic parameters to real timescales for both approaches requires a mutation rate and generation time. Our chosen value of 5 years for generation time is widely used for *Acropora* (Mao et al. 2018; Matz et al. 2018; Cooke et al. 2020) and reflects its fast growth rate combined with high mechanical vulnerability of older colonies (Madin et al. 2014). For the mutation rate we calculated an updated value (µ=1.2e^−8^) based on recently published divergence times (Shinzato et al. 2020). To capture uncertainty in both parameters we ran demographic analyses with SMC++ using alternative published values for the mutation rate (µ=1.86e^−8^, 2.98e^−8^) and alternative plausible values for generation time (3y, 7y). Variation in these parameters did not result in qualitative changes to the shape of Ne curves, but generally led to more-recent estimates for key events such as bottlenecks and population splits (supplementary fig. S10).

Changes in effective population size (Ne) during the past 1My inferred by SMC++ revealed qualitatively similar trajectories for the three populations identified in population structure analyses. All experienced a strong bottleneck some time between 7 and 15 Kya followed by expansion and stabilisation. Timing of these bottlenecks coincides with a period of rapid sea level rise at the end of the last glacial maximum (fig 2B). In agreement with the existence of a bottleneck and subsequent population expansion, genome-wide estimates of Tajima’s D for all three populations were negative (supplementary fig. S11).

**Fig. 2.**
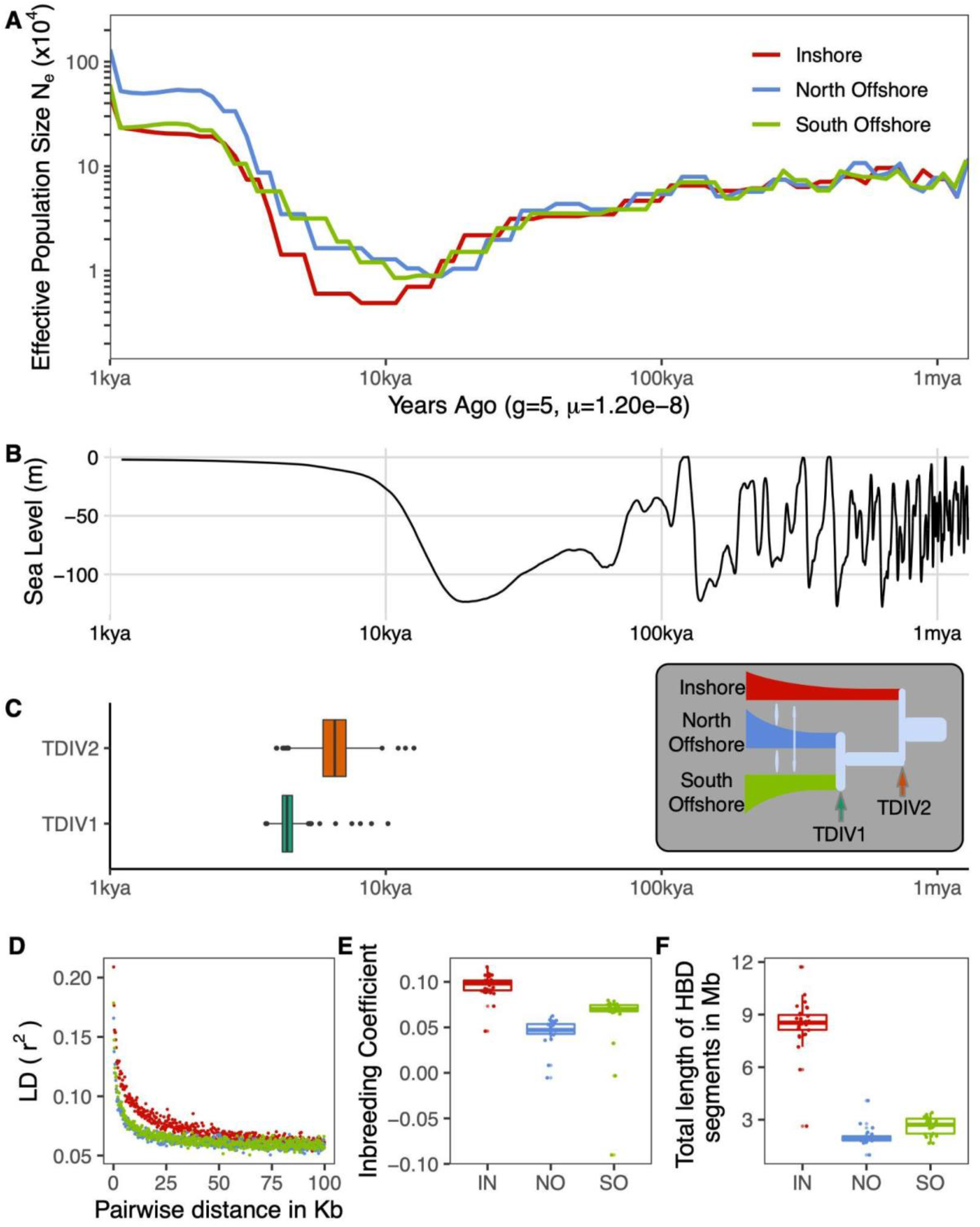
Demographic history of *A. digitifera* in Western Australia during the past 1 million years. Locations are denoted by two letter codes, inshore (IN), north offshore (NO), south offshore (SO) and coloured as shown in A. **A**. Changes in effective population size (Ne) inferred by SMC++. **B**. Change in global sea level over the same timescale as depicted in A (data from Bintanja and Wal 2008). **C**. Estimated divergence times for the inshore-offshore split (TDIV2) and offshore split (TDIV1) obtained using fastsimcoal2. Inset shows the best model; also used to fit bootstrap parameter estimates. **D**. linkage disequilibrium (LD) decay calculated using plink. **E**. Boxplot of the inbreeding coefficient calculated using plink2 for each sample. **F**. Total length of genomic regions within each individual that were homozygous by descent (HBD) calculated using ibdseq (Brian L. Browning and Browning 2013). All demographic parameter estimates for both SMC++ and fastsimcoal2 were scaled to real times based on a generation time of five years and an estimated mutation rate of 1.2×10^−8^ per base per generation.

Populations differed in the timing and severity of the bottleneck, with the strongest and most recent effects seen inshore. This was evident in the SMC++ trajectory as well as the much higher prevalence of homozygous-by-descent (HBD) segments in inshore (fig. 2F) along with elevated inbreeding coefficients (fig 2E) and linkage disequilibrium (fig. 2D). Differences between the two offshore populations were less pronounced than between offshore and inshore, however it was clear that the north offshore population retained the highest overall levels of diversity as it had the lowest inbreeding coefficient, smallest proportion of HBD segments and highest SMC++ estimated Ne during the recent stable period (2-5Kya).

Divergence time estimates from both SMC++ and fastsimcoal2 indicate a recent split for all three populations that coincides with the same post-glacial time window as bottlenecks observed in SMC++ analyses. Bootstrap estimates for the inshore-offshore split based on the best-fitting model in fastsimcoal2 (fig. 2C; supplementary table S8B) were older (5-8Kya) than those between offshore locations (4-5Kya), matching our expectations based on pairwise Fst values and population structure analyses (see above). Estimates from SMC++ were in approximate agreement with this (9Kya) but did not differentiate between inshore-offshore and offshore-offshore splits.

In addition to estimating split times, we used fastsimcoal2 to test a range of competing demographic scenarios (supplementary fig. S13). The results indicate that a model IMc (fig. 2C inset) with constant migration between offshore populations and secondary contact between inshore and offshore provides a better fit to the SFS than competing models with strict isolation (SI), ancient migration (AM) or continuous migration (IM) (supplementary table S8B). Support for a model (IMc) with contemporary migration was surprising given the lack of evidence for gene flow in admixture analyses but is reconciled by the fact that estimated migration rates from the IMc model were extremely low (~1e-4) (supplementary table S8B). To confirm that the IMc model is consistent with this and other key features of our data we calculated summary statistics and performed admixture analyses for simulated data under this model. These analyses (summarised in supplementary fig. S15) showed similar patterns of HBD, inbreeding coefficient and admixture to our results based on sequencing (fig 1), but produced positive values for Tajima’s D (negative in our real data). Tajima’s D is sensitive to the recency and strength of bottlenecks and a discrepancy between modelled and observed values suggests that the IMc model may be underestimating their age or strength (Gattepaille et al. 2013).

Strong bottlenecks and low migration are both potential contributors to population differentiation. To estimate the relative contribution from these factors, we ran simulations based on the IMc model, but with bottlenecks removed by setting a constant effective population size (equal to the ancestral value) and other parameters, including split times and migration rates set to their best-fitted values. Compared with simulations under the full model, removing the bottleneck dramatically reduced pairwise Fst by five fold for the inshore-offshore split and 2.5 fold for the split between offshore locations (supplementary fig. S16A).

### Genome-wide scan for selective sweeps

To investigate the effects of natural selection on the *A. digitifera* populations identified above we performed a genome-wide scan for signatures of selective sweeps (regions of low diversity arising due to positive selection and linkage to a beneficial allele). As the primary basis for this scan, we used three statistics (iHS, XP-EHH, XP-nSL) that summarise patterns of extended haplotype homozygosity (EHH) because these have high power to detect selective sweeps within independent populations (iHS) (Voight et al. 2006) or as a contrast between pairs (XP-EHH;XP-nSL) (Sabeti et al. 2007; Szpiech et al. 2021). Following standard binning and normalisation practice (see methods; Szpiech and Hernandez 2014) we identified a total of 231 loci (50kb windows) in which at least one of these three statistics was significant (top 1%) based on the frequency of occurrence of SNPs with extreme values. These putative sweep loci were spread throughout the genome (fig 3A; supplementary table S4) and included 72 specific to inshore, 80 to south offshore, and 79 to north offshore. They were also enriched in SNPs for which the allele-frequency-based indicator of selection, population branch statistic (PBS), had extremely high values (fig 3A).

**Fig. 3.**
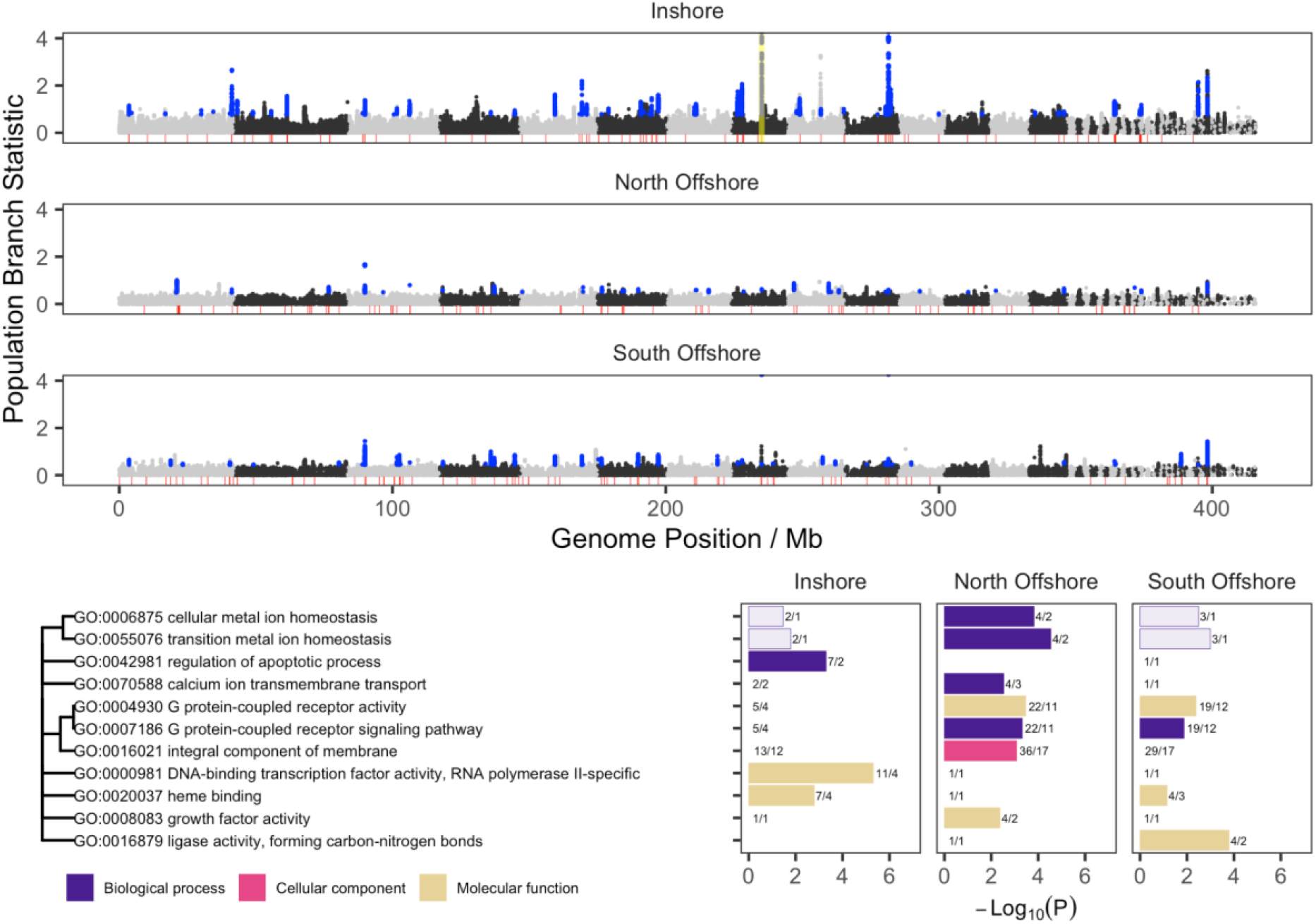
Genome wide distribution of signatures of selection and functional enrichment for overlapping genes. **A**. Manhattan plots showing values of the population branch statistic (PBS) and regions under selection identified by EHH based scans. PBS estimates are shown as points for each population with the other two considered as outgroups. Points are shown in black and grey to indicate transitions between alternating pseudo-chromosomes via mapping to the *A. millepora* assembly from (Fuller et al. 2020). The red shaded baseline shows the location of regions identified as candidates for positive selection using EHH-based scans. Blue points indicate PBS values with probability of false discovery less than 1% under the best fitting demographic model, and which are coincident with EHH scans. Yellow highlighted region in Inshore shows the location of the peroxinectin locus. **B**. GO term enrichment for regions under selection in inshore and offshore populations. Bar colour indicates one of three broad ontologies, BP: Biological Process, CC: Cellular Compartment, and MF: Molecular Function. Relationships between enriched terms based on numbers of shared genes are shown as a dendrogram (left). Length of bar indicates the log odds of enrichment (−Log10(p)) based on p-values calculated from Fisher’s exact test. Numerical labels indicate the number of genes putatively under selection followed by the number of loci intersected by those genes. Dark shaded bars show significant enrichment based on numbers of genes and numbers of independent sweeps while light shaded bars are significant based on numbers of genes but not sweeps.

To control for demographic effects such as bottlenecks we used simulated data under the best-fitting (IMc) demographic model to calculate threshold values for the PBS that would result in fewer than 1% false positives. As expected, given its more severe bottleneck, this threshold was higher for inshore (0.76) compared with offshore populations (NO:0.48, SO:0.44). Even at this higher threshold however the inshore population had more sweep regions identified by EHH statistics that also overlapped SNPs with significant PBS values (33/72, 45%) compared with north offshore (18/79, 23%) and south offshore (25/80, 31%).

Of the 1015 genes that overlapped with loci putatively under selection (231 loci identified via EHH-stats; see above), 515 could be assigned a GO term using InterProScan 5 (Jones et al. 2014) based on gene family membership inferred from the presence of conserved domains. Analysis with topGO revealed a total of 11 GO terms across all three ontologies (6 MF;5 BP; 1 CC) that were enriched (p<0.005; at least 2 distinct sweep regions) in these genes (supplementary table S5) compared with the background (supplementary table S9) in one or more of the three populations (fig. 3B). Since multiple genes often overlapped with each sweep region, we also calculated enrichment statistics based on sweep regions rather than genes as independent units, and found that all these terms were also enriched (Fisher’s exact test p<0.005) in at least one population under this criterion (fig. 3B).

Three groups of GO terms showed exclusive enrichment in either inshore or offshore locations, potentially reflecting broad patterns of selection related to contrasting environmental conditions. Terms related to membrane G protein-coupled receptors (GPCRs) (GO:0004930, GO:0007186, GO:0016021) were strongly enriched in both offshore populations but not in the inshore, with genes underpinning this pattern distributed across 23 independent sweep regions. Exclusive enrichment in inshore was observed for the GO terms, transcription factor activity (GO:0000981) and regulation of apoptotic process (GO: 0042981). Genes supporting enrichment of transcription factor activity in inshore included a diverse range of transcription factors including those containing homeobox, C2H2 zinc finger, T-box, and fork head domains, all of which are involved in regulating early development. Enrichment for the GO term, apoptotic process was supported by two independent sweeps, one containing a Bcl-2-like protein (IPR026298) and another that hosted a cluster of 6 genes each containing a single death effector domain (IPR001875).

### Selective sweep at the peroxinectin locus

To investigate the link between selection, climate change, and gene function in additional detail we chose to focus on one of the strongest signatures of selection in the inshore population. This locus was associated with the highest PBS values (yellow highlight in fig 3A), low Tajima’s D (fig 4A), and had a clear differentiation between selected and background haplotypes (fig 4A). It also contained by far the largest number (84; next-highest, 7) of near privately fixed SNPs (>90% allele frequency in inshore, absent in offshore), and of these, over 90% were contained within a single gene, s0150.g24.

**Fig. 4.**
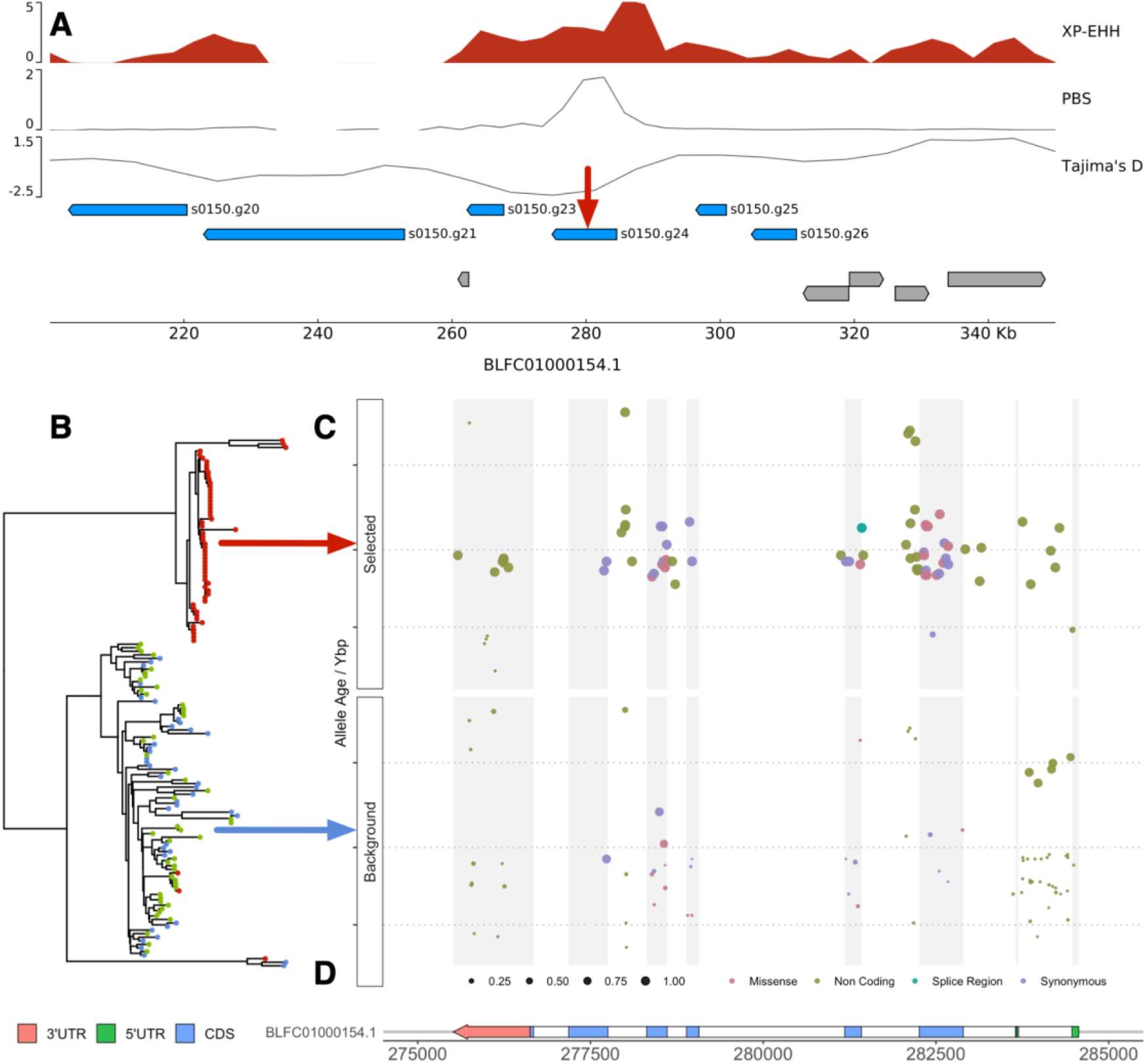
Gene arrangement, haplotype structure and timing of selection for a selective sweep at the peroxinectin locus. **A**. Zoomed detail at the locus highlighted in yellow in fig 3A. Tracks show values for XP-EHH, PBS and Tajima’s-D for the inshore population. Horizontal bars show the location of genes with peroxinectins in blue and all other genes in grey. **B**. Neighbour joining tree (left) based on core haplotypes. Core haplotypes include 200 phased variant sites centred on position 281245 on scaffold BLFC01000154.1 (shown with a red arrow in A). Each haplotype is shown as a terminal branch in the tree and coloured according to sample location. Haplotypes with the derived allele at the focal SNP all partition into the top clade (selected haplotypes) and those with the ancestral allele into the bottom clade (background). **C**. Age, consequence and frequency of variants overlapping the gene s0150.g24. Scatterplots show variants on selected haplotypes (top) and background haplotypes (bottom). Point positions reflect genomic coordinate (x-axis) and age (y-axis). Points are coloured according to their variant effect category inferred by BCFtools csq and sized by allele frequency. Allele frequencies are calculated as the proportion of haplotypes with the derived allele in the given population grouping, i.e selected or background. Grey vertical bars serve as guides to indicate the position of exons. **D**. Gene structure of s0150.g24 showing exons, cds and untranslated regions.

Unlike many other sweep loci where the diversity of genes makes it difficult to associate gene function with selection, four of the five genes overlapping this 50kb sweep region encoded peroxinectin-like proteins (Panther subfamily PTHR11475:SF4; CDD cd09823) and these formed part of a cluster of 8 peroxinectin genes found within 200kb of the sweep. A genome-wide search for haem peroxidases (IPR019791), the parent superfamily that contains peroxinectins, revealed a total of 15 in *A. digitifera*, however only one additional peroxinectin-like gene was found outside the peroxinectin locus. All remaining haem peroxidases were scattered on different scaffolds throughout the genome indicating that peroxinectins, but not haem peroxidases in general are co-located. Orthologous genomic clusters of peroxinectins were also present in other *Acropora* species (*A. millepora, A. tenuis*; supplementary fig. S17) indicating that the arrangement is at least as old as the crown age of this genus (~50Mya; Shinzato et al. 2021).

The strongest statistical indicators of selection at the peroxinectin locus are centred on the gene, s0150.g24 (fig 4A). An estimate for the timing of selection on this gene based on the inferred time to the most recent common ancestor for selected haplotypes (8.0-8.3Kya; starTMRCA Smith et al. 2018) approximately matches the divergence time for inshore corals. It is also consistent with the relationship between frequency and age of individual alleles at SNPs in this gene inferred by GEVA (Albers and McVean 2020). Young alleles (aged less than 8 Kya) had low frequencies in both selected and background haplotypes, consistent with their emergence after the sweep, whereas alleles older than 8Kya showed a strong shift toward high frequencies in selected haplotypes compared with background (fig 4C; supplementary fig. S18).

Examination of the consequences of variants within the gene, s0150.g24 suggests that selected haplotypes may encode a change in exon usage. We identified a total of 10 missense variants in the third exon in selected haplotypes compared with just one at low frequency in the background. Such accumulation of variation in an otherwise conserved region suggests that this exon may no longer be expressed. Although more work is required to confirm this we note that several variants that might encode the change are present, including a change in the splice region between the third intron and fourth exon as well as five variants in the first intron, a region that often contains gene regulatory elements (Chorev and Carmel 2012).

## Discussion

Our results demonstrate rapid divergence in *A. digitifera* from northwestern Australia resulting in three genetically distinct populations separated by location. Estimated split times of 5-10Kya and similarly timed bottlenecks in all three populations coincide with geological evidence for the post-glacial reestablishment of reef growth on the tops of atolls (Collins et al. 2011) and inshore reefs (Solihuddin, Bufarale, et al. 2016) in this region. Simulations based on our best fitting demographic model showed that population size changes were a major contributor to overall levels of population differentiation, most likely through increased genetic drift at small population sizes. Limited dispersal indicates that these bottlenecks are likely to represent founder effects arising from post-glacial colonisation, and the two factors (low dispersal and bottlenecks) are the main neutral drivers of divergence.

Since many marine taxa have pelagic larvae and large species ranges it was initially thought that they should exhibit limited or weak population structure (Palumbi 1992; Cowen and Sponaugle 2009). Recent advances in our understanding of larval dispersal in corals and reef fishes have shown that both can be highly variable (Jones et al. 2009), indicating that in suitable settings strong population structure may be present (Underwood et al. 2020). In agreement with this, population structure has now been observed for a range of coral reef taxa (Warner et al. 2015; Lukoschek et al. 2016; Underwood et al. 2018; Thomas et al. 2020) but the mechanisms giving rise to this diversity remain poorly understood. Our study demonstrates that population structure can arise rapidly (<10Kya) when dispersal is low, especially if this is combined with colonisation of new habitats thereby inducing founder effects which enhance drift. Strong selection (as observed in our study) might also contribute to population structure, however, our neutral simulations show that this is not required to account for rapid divergence.

The limited connectivity inferred between locations in northwestern Australia is in contrast to studies of other acroporid species on the Great Barrier Reef (GBR) (Lukoschek et al. 2016; Cooke et al. 2020; Fuller et al. 2020), and the Ryukyu Archipelago (Shinzato et al. 2015). Both *A. tenuis* and *A. millepora* on the GBR form near panmictic populations with weak isolation by distance structure over hundreds of kilometres along north-south stretches of the reef (Lukoschek et al. 2016). Where highly differentiated populations do exist (eg *A. tenuis*; Cooke et al. 2020) they show signs of recent admixture and likely reflect ancient splits that are now in secondary contact. This high level of connectivity most likely reflects the fact that reefs in the GBR form a continuous chain with spacing between 1 and 50km (Almany et al. 2009), and those in the Ryukyu are connected by the Kuroshio current (Shinzato et al. 2015). In contrast, reefs in Western Australia are relatively isolated on offshore atolls or inshore islands separated by distances of 100’s of km (Wilson 2013). The results of this study therefore highlight the potential for physical distances combined with a lack of intermediate habitats to act as a barrier to gene flow, even in a broadcast spawning marine species with a pelagic larval stage. It also underscores the importance of historical context and demographic modelling when interpreting measures of genetic differentiation such as Fst. In this case low Fst did not mean high connectivity as in Wright’s Island model (Wright 1931), but recent divergence.

### Contrasting selection between inshore and offshore habitats

We identified clear evidence for selection across a wide diversity of loci in all three populations, but with the strongest signals observed in the inshore. The inshore reefs of northwestern Australia are notable for their extreme temperatures (short term maxima of 37 °C), frequent aerial exposure at low tide and highly variable turbidity (Wilson 2013; Solihuddin et al. 2015). The complex, polygenic nature of these stressors, combined with the fact that signatures of selection often cover many genes (due to linkage) make it difficult to identify causal alleles or genes. These issues affect most previous work attempting to identify the genetic basis for heat adaptation in corals (Dixon et al. 2015; Fuller et al. 2020; Smith et al. 2022). In our study, however, we identified a highly localised signal on a gene (s0150.g24) within a locus dominated by other genes from the same family (peroxinectin-like haem peroxidases).

Peroxinectins are best characterised in arthropods where they mediate the immune response via cell adhesion (Johansson et al. 1995) and prostaglandin synthesis (Park et al. 2014). Heat stress experiments in molluscs (Lang et al. 2009), and corals (Voolstra et al. 2009; Shinzato et al. 2021; Traylor-Knowles et al. 2021) consistently identify peroxinectin-like proteins as differentially expressed, and there is evidence that they have undergone recent expansion in some heat-tolerant coral lineages (Shinzato et al. 2021). Unfortunately the role of peroxinectins in corals has been obscured because many peroxinectin-like proteins are annotated as peroxidasin homologues in the NCBI nr database. For three key publications (Voolstra et al. 2009; Shinzato et al. 2021; Traylor-Knowles et al. 2021) we manually checked sequences annotated as peroxidasin-like, and that were differentially expressed in response to heat stress and found that in all cases the corresponding protein sequences had a similar domain structure to the peroxinectins identified in this paper. All contained one or more characteristic conserved domains of peroxinectins (Panther subfamily PTHR11475:SF4 or CDD cd09823) but lacked the N-terminal leucine rich repeats and immunoglobulin domains found in peroxidasins.

Our results highlight the potential importance of peroxinectins in adaptation to the extreme conditions experienced by inshore corals, and invite future work to characterise the evolution and function of co-located peroxinectins in Acropora and related taxa. In particular, since the selected haplotypes differ in amino acid sequence to the background, further functional genetic work has a strong chance of identifying the precise nature of the beneficial change, thereby providing a rare opportunity to associate gene function with local adaptive benefit in a wild population.

### Implications for conservation

Our results document dynamic population responses of *Acropora digitifera* to past climate change. They suggest that this species was likely extirpated throughout much of Western Australia during the last glacial maximum, but recolonised and underwent rapid population expansion when conditions became favourable. This history may contribute to their vulnerability; particularly for the inshore populations of the Kimberley, which despite their remarkable ability to survive extreme heat, turbidity and exposure, have suffered a loss of genetic diversity due to founder effects and strong selection.

We also observed low natural connectivity between inshore and offshore sites. Connectivity (and gene flow) in coral populations is a key deciding factor in their ability to adapt to climate change (Matz et al. 2018) because it allows natural selection to act on a larger overall gene pool, and because it mitigates against local losses.

This combination of risk factors (bottlenecks and low connectivity), seen in our study may also be present in other coral reef systems with similar biogeography such as widely spaced offshore atolls and island chains. Our results therefore suggest that corals from northwestern Australia and other similar systems may be at a higher risk from climate-related losses than in highly connected systems such as the Great Barrier Reef.

## Materials and Methods

### Sampling and sequencing

75 samples from adult corals across our three study locations were selected from a larger pool of 564 samples collected as part of a separate study across a wider geographic area that was primarily based on DArT sequencing (Adam et al., under review). At these three study locations, small nubbins of *A. digitifera*, approximately 1-6 cm3 were collected in November 2017 (Rowley Shoals, Ashmore Reef, Adele Island and Beagle Reef) and March 2018 (Rowley Shoals) across 21 sites and stored in 100% ethanol. These samples were later subsampled to be sent to Diversity Array Technology Pty Ltd. (DArT P/L) for further processing. DNA extractions were performed for all samples by DArT and the remaining DNA (not used for DArT) was sent to the QB3 UC Berkeley sequencing centre for whole genome sequencing. Samples for whole genome sequencing were selected randomly from samples previously sequenced by DArT and after excluding 7 that failed initial quality checks an additional 7 replacement samples were also randomly selected. Initial sequencing was performed on a single NovaSeq S4 flowcell to obtain ~3 billion 2×150bp paired-end reads across all samples. Additional sequencing was then performed on a second NovaSeq S4 flowcell for 33 samples because they failed to achieve the target depth of 10x in the first batch. Samples included in the second batch of sequencing were spread across all sites in the study (supplementary table S1) and we did not observe any population structure attributable to batch in fineSTRUCTURE analyses (supplementary fig. S3).

Initial population structure analyses revealed a single sample (BR_5_121_S125) coded as inshore that clustered with south offshore samples. To check whether this was a genuine example of migration or a mislabelled sample we combined our whole genome sequencing data with all raw reads from the DArT dataset. Raw DArT reads were first mapped to the genome using bwa mem (v0.7.17) and variants called with freebayes (v1.3.2-dirty; Garrison and Marth 2012) with min-mapping-quality set to 30 and min-base-quality set to 20. The resulting vcf file was then filtered to retain only variants with maf>0.1, min depth of 8x and min mean depth of 15. This vcf was then merged with the filtered vcf file from whole genome analyses retaining only variant sites common to both approaches. Using this combined vcf we then calculated the relatedness using the relatedness2 statistic implemented in VCFtools (v0.1.16) between all pairs of samples and found that all but two pairs had relatedness values < 0.1. The remaining two pairs had relatedness values (>0.48) indicative of clones or identical samples. One of these pairs was irrelevant to the current analyses as it concerned two DArT samples only. The remaining pair indicated a match between sample RS3_S_252 from the DArT dataset and BR_5_121_S125 from the WGS dataset indicating that this sample was mislabelled at some point after DArT sequencing, and its true origin was the Rowley Shoals.

### Data pre-processing and variant calling

Sequencing reads for all samples were checked for quality issues using FastQC version 0.11.9 and multiQC version 1.6 (Ewels et al. 2016). No samples were flagged as poor quality on the basis of these checks. We then followed the GATK4 (4.1.9) (McKenna et al. 2010) best practice workflow for germline variant calling. Key workflow steps were as follows; raw reads were first aligned to the *Acropora digitifera* reference genome (Shinzato et al. 2011; Shinzato et al. 2020) using BWA version 0.7.17 (Li and Durbin 2009) with the BWA-MEM algorithm; duplicated reads were removed using the MarkDuplicates function in GATK. Next, HaplotypeCaller was used to call variants in each dataset and generate a file in the GVCF format. The GVCFs from all samples were consolidated into a GenomicsDB datastore using GenomicsDBImport and passed to the joint genotyping tools GenotypeGVCFs. The whole workflow was implemented using snakemake version 5.5.4 (Koster and Rahmann 2012) and is available online at (https://github.com/bakeronit/snakemake-gatk4-non-model).

The initial variant call set was filtered with the objective of minimising bias while maintaining quality biallelic SNPs suitable for the population genomic analysis. Filtering steps were performed sequentially as follows;

1. Sites within 5bp of InDels were removed using BCFtools version (1.10.2) (Danecek et al. 2021)
2. Hard-filtering thresholds were applied using the GATK VariantFiltration tool based on recommended parameters as follows (QD < 10, QUAL < 30, SOR > 3, FS > 60, MQ < 40, MQRankSum < −12.5, ReadPosRankSum < −8). Abbreviated parameters are QD=QualByDepth, QUAL=Quality, SOR=StrandOddsRatio, FS=FisherStrand, MQ=RMSMappingQuality.
3. Sites located in simple repeat regions identified by mdust version 2006.10.17 were removed (Li 2014).
6. Sites were removed if they had more than 10% missing or low quality genotype calls under the thresholds GQ > 20 and DP >3. (GQ=Genotype Quality; DP = sample read depth). This was performed using VCFtools v0.1.16 (Danecek et al. 2011)
7. Sites were removed if their read coverage fell outside expected bounds (mean per-sample depth less than 8 or greater than 31) because this could indicate collapsed repeats or regions with low mappability.

After all filtering steps, we obtained 9,656,554 high-quality biallelic SNPs from 75 samples. A summary of the number of missing genotypes in all samples after filtering is provided in supplementary fig. S1B.

### Phasing and imputation

To resolve haplotype information, the read aware phasing mode of SHAPEIT v2 (Delaneau et al. 2012) was used to phase all segregating sites in the filtered VCF file for every scaffold. Phase informative reads in mapping files were first extracted by extractPIRs. Next SHAPEIT assembled these reads into haplotypes and converted them into VCF format.

Missing genotypes were imputed by SHAPEIT2 during the assembly run. To evaluate the accuracy of imputation, we performed a “masked analysis” (Verma et al. 2014), in which a subset of genotyped SNPs in the samples was randomly pruned and then imputed as missing data. We compared the imputed genotypes to their original genotypes to estimate the concordance which indicates the performance of imputation with respect to that set of SNPs. A summary of this imputation accuracy check is provided in supplementary fig. S2.

### Genome-wide population genetic statistics

Nucleotide diversity(*π*), Tajima’s D, linkage disequilibrium, and heterozygosity were calculated genome-wide using the unphased, filtered variant set. The het function in PLINK2 (v2.00a3) (Chang et al. 2015) was used to calculate heterozygosity in each sample. Nucleotide diversity and Tajima’s D were both calculated in 10kb windows with a 2kb overlap using VCFtools and VCF-kit (Cook and Andersen 2017) respectively. To avoid bias from gaps and masked regions in these window-based estimates, we used BEDTools v2.29.2 (Quinlan and Hall 2010) to remove windows that have less than 70% of bases covered, leaving 136,435 windows. Pairwise linkage disequilibrium (r^2^) was calculated in 1Mb windows using plink v1.9 (Purcell et al. 2007) based on an equal number (20) of samples from each location. Pairwise Fst for all SNPs was calculated using the weir-fst-pop function in VCFtools.

### Population structure

We performed population structure analysis using complementary methods based on allele frequencies (PCA, ADMIXTURE) and phased haplotypes (fineSTRUCTURE, IBD sharing). For the allele-frequency-based analyses we started with the unphased, filtered variant set and then performed further filtering to remove sites with minor allele count less than or equal to one, and that deviated significantly from Hardy-Weinberg equilibrium (p-value < 1e^−4^). To minimize the effect of SNPs in high linkage disequilibrium (LD) for PCA and admixture analysis, we filtered out SNPs based on pairwise LD using PLINK v1.9 (—indep -pairwise 50 10 0.1). PCA analysis was performed using smartpca from EIGENSOFT v6.1.4 (Price et al. 2006) with LD pruned SNPs. Admixture analysis was performed on the same LD pruned data as the PCA analysis using ADMIXTURE v1.3.0 (Alexander et al. 2009). ADMIXTURE was run using 10-fold cross validation and the number of clusters was varied from 1 through to 6. Although the cross-validation error was lowest for K=1, we chose to use K=3 because it reflected the number of clusters seen in PCA and because inference of K=1 is common in situations where overall divergence between clusters is low (Lawson et al. 2012).

As a complement to the allele-frequency-based analyses of population structure we also performed a fineSTRUCTURE (version 4.1.0) analysis (Lawson et al. 2012) on the phased dataset. As input to this analysis we used the phase files generated by SHAPEIT (converted with impute2chromopainter.pl) and a recombination map file that was generated assuming a uniform genome-wide recombination rate. We used the linked mode to take advantage of phasing information in our data and allowed the Markov Chain Monte-Carlo (MCMC) to run for 2,000,000 iterations with a burn-in of 1,000,000. Tree inference was performed with 10,000 maximization steps.

To identify patterns of relatedness between samples based on genomic regions inherited by descent (IBD) we first identified IBD segments using the package Refined IBD (Brian L. Browning and Browning 2013). Breaks and short gaps in segments were removed using the companion program merge-ibd-segments. Pairwise relatedness was calculated using a python script (relatedness_v1) provided by the authors of Refined IBD (Browning and Browning 2011) based on the total length of shared haplotypes as a proportion of total genome size.

### Population structure including Japanese samples

We used PCAngsd (Meisner and Albrechtsen 2018) to perform a principal components analysis (PCA) of our Western Australian samples together with publicly available data on 16 samples of *A. digitifera* from Japan (Shinzato et al; Bioproject PRJDB4188). Raw reads from Japanese samples were mapped to the *A. digitifera* genome using BWA MEM and duplicates marked using GATK as was done for our own samples. We then used ANGSD (v 0.933) (Korneliussen et al. 2014) to call and filter SNPs as well as calculate genotype likelihoods based on data from all bams from Western Australia and Japan. ANGSD was run with the GATK genotype likelihood mode and SNPs were filtered to remove sites with data on fewer than 90% of samples, with quality scores less than 20, overall read depth less than 910 or greater than 3000, minor allele frequency < 0.05 or with a Hardy-Weinberg p-value less than 1e^−6^. This resulted in a beagle formatted genotype likelihood file which was used as input to PCAngsd (v0.98) that was run with default settings. The PCA shown in supplementary fig. S5 was produced based on the resulting covariance matrix.

The realSFS program included as part of ANGSD was used to calculate Fst values between all pairs of populations (shown in supplementary table S3B). This was done only when comparing with Japanese samples since it was not possible to use plink (used elsewhere to calculate Fst) when combining data with highly variable depth of coverage. First ANGSD was used to export allele frequencies at the same sites used for PCAngsd (see above) and these were then used as input to realSFS to calculate 2D sfs files for each pair of populations and then to calculate Fst values.

### Phylogenetic inference based on UCE and Exon probes

To place the *A. digitifera* populations from this study within a broader phylogenetic context we extracted established phylogenetic markers (ultra-conserved-element and exon sequences from Cowman et al. 2020) from our Western Australian samples, previously published data from Japanese samples (Shinzato et al. 2015), and published reference genomes for *Acropora millepora* (Ying et al. 2019) *and Acropora tenuis* (Cooke et al. 2020). For this analysis we used a randomly selected subset of three samples from each of our populations and three from Japan. First we mapped the hexa-v2 probeset (Cowman et al. 2020) to the genomes of all three species (*A. digitifera, A. tenuis, A. millepora*) using BWA (v0.7.17). We then used BCFtools (1.11) to call a consensus sequence corresponding to a 1000bp interval around the central base of each probe. Ambiguous bases arising from heterozygous sites were encoded using their corresponding IUPAC ambiguity codes in this process. BEDTools (v2.30.0) was used to merge overlapping intervals and extract the called consensus sequences in fasta format. We then used MAFFT (v7.394) (Katoh et al. 2002) to align sequences for each (~1000bp) region separately. We then created a partition file in Nexus format listing alignments for the 1681 sequences present in all samples. Finally we ran IQ-TREE (v2.0.3; Nguyen et al. 2015) with the option MF+MERGE which uses modelfinder (Kalyaanamoorthy et al. 2017) to identify the best model for each partition and then used this optimised partition scheme to build a tree with 1000 ultrafast bootstraps (Hoang et al. 2018). The resulting tree (supplementary fig. S7) placed the *A. digitifera* genome sequences together with all the Western Australian and Japanese *A. digitifera* population samples in the same clade, but was unable to resolve the population-level differences identified through allele-frequency analyses (above). Branch lengths within this *A. digitifera* clade were very short relative to known species-level relationships such as those between *A. digitifera, A. tenuis* and *A. millepora* (supplementary fig. S7). Given that UCEs have recently been shown to resolve Acropora species (Cowman et al. 2020), these results confirm that all of the distinct populations identified in the present study are likely to be conspecifics.

### Demographic history with SMC++

SMC++ analysis was performed based on the unphased vcf callset after having removed one mislabelled sample (BR_5_121_S125; see section on sequencing and sampling). To avoid problems arising from highly fragmented scaffolds only those with a length greater than N90 (107,903bp) were used. The vcf files of each scaffold were converted into SMC++ input format with large uncalled regions masked using the vcf2smc script. To perform composite likelihood estimates, multiple SMC files were generated for each scaffold by varying the choice of “distinguished individual” over the same VCF to every individual (inshore: 29, north offshore: 20, south offshore: 25). To estimate population size histories, all SMC++ input files were used together in a single run with the options “thinning 3000, 50 EM iterations, 40 knots, with mutation rate same as before (1.20e^−8^), setting the starting and ending time points to 20-200000 generations”. The divergence times of each population pair were also inferred using the SMC++ split command with marginal estimates produced by using the estimate option.

To address the uncertainty in SMC++ analysis from mutation rate and generation time parameters, we applied the same estimate run with the same parameters except mutation rate which we tested two other alternative mutation rates: 1.86e^−8^ (Cooke et al. 2020); 2.98e^−8^ (Mao et al. 2018) for three populations. For each run, we generated the effective population size curve with three generation time 3, 5, and 7 years (Oppen et al. 2000; Baria et al. 2012; Matz et al. 2018). We also used results generated with different mutation rates in splitting time estimate.

### Demographic history with fastsimcoal2

To model demographic history while accounting for population structure, we carried out SFS based demographic modelling using fastsimcoal2 (Excofffier et al. 2021). We used all samples except BR_5_121_S125 as per our SMC++ analysis. To minimise the bias from linkage disequilibrium and selection, we used BCFtools to remove sites located in genic regions and performed LD pruning in 1000bp windows with a cut-off of r^2^>0.3. To utilise the mutation rate in branch length computation, we estimated the monomorphic sites based on the proportional number of mappability sites defined by the SNPable pipeline we used in MSMC analysis. We also filtered out sites with missing genotypes and then used easySFS (https://github.com/isaacovercast/easySFS) to generate a joint three-dimensional folded SFS with 257,314 SNPs.

We firstly tried to test which population tree topology the SFS data support without considering the population size changes and migrations. In this step we tested four alternative topologies indicating alternative splitting modes among three populations including inshore split first, south offshore split first, north offshore split first, or a polytomy tree of three populations (supplementary table S8A). For each model, fastsimcoal2 (version 2705) was used to fit parameters to the joint SFS with 50 ECM optimization cycles and 200,000 coalescent simulations used to compute the likelihood. This model fitting process was repeated 100 times based on different randomly sampled starting parameter values. This gave clear support for the inshore split first model as it always had the lowest Akaike information criterion (AIC) value across all 100 runs. We report the best AIC and likelihood values for all four models (across the 100 runs) in supplementary table S8A.

Based on the population tree ((NO, SO), IN), we then tested six competing models all with exponential population size change (supplementary fig. S12). These models were primarily designed to test different migration scenarios and comprised; 1) strict isolation (SI), 2) continuous migration between all demes at all times (IM), 3) continuous migration among three populations only after offshore divergence, ie secondary contact for offshore-inshore but isolation with migration for offshore-offshore (IMc), 4) isolation with recent secondary contact (SC), 5) early migration after offshore divergence (EM), 6) ancient migration between inshore and offshore ancestor followed by strict isolation (AM). We specified the search ranges for the current and ancestral effective population sizes between 1,000 and 1,000,000, and the effective population size for the offshore ancestor to between 100 and 10,000, but with an open upper bound that is extended if parameters get close to the boundary during the ECM optimisation. Divergence times were allowed to vary between 100 and 10,000 generations. The range of migration rates was assumed to be between 10^−7^ to 10^−3^ with open upper bounds. For the SC and EM models, we allowed the time of changed migration (TMIG) to be between 100 generations and the offshore divergence time (using paramInRange).

Parameters for all six models were initially estimated using the same process as outlined above. After parameter estimation, we observed that the SC and EM models were converging towards the IMc model as TMIG kept being pushed to the lower bound (100 generations) in the EM model while being optimised to be close to offshore divergence time in the SC model (supplementary fig. S13). We thus deprecated these two models in the following likelihood estimates and model comparison. Next, we compared different models using the model normalised relative likelihood (Excoffier et al. 2013) (supplementary fig. S14, supplementary table S8B), and estimated the parameter ranges (supplementary table S8B). As a result of this process we chose the IMc model as it had a model normalised relative likelihood of close to 1 whereas this was 0 for all other models. We then estimated confidence intervals for the parameters of the best model using 100 non-parametric bootstrapping datasets, each of which was generated by sampling 257,314 SNPs with replacement from the original set of SNPs. This sampling was performed using the sample tool (alexpreynolds.github.io/sample). For each bootstrapping data set, we performed 20 independent runs. Final results shown in supplementary table S8B show 95% confidence intervals based on the distribution of fitted parameters from these independent runs.

### Analysis of simulated data under the best fitting model

To verify that our best-fitting demographic model obtained with fastsimcoal2 (model IMc) was not only a good fit to the SFS, but also to other population genetic parameters we generated simulated data under this model. Simulations were performed using fastsimcoal2 using an identical model specification file to that used for SFS fitting. We performed 50 independent simulations, each of which used parameters drawn randomly from a uniform distribution across a 90% confidence interval based on our bootstrap estimates (see above). Each simulation generated 20 scaffolds of length 2mb with the goal of mirroring the approximate fragmentation level of our real genome.

Based on this data we then calculated; (1) the length of HBD segments using ibdseq, (2) inbreeding coefficient using plink2, (3) Tajima’s D using vk tajima, (4) admixture coefficients using ADMIXTURE, (5) population branch statistics using plink. All calculations were performed using identical settings to those used for real data. The results are shown in supplementary fig. S15.

Simulations based on a modified version of the IMc model were used to assess the contribution from population size changes (ie the bottleneck) to population differentiation. The IMc model was modified so that the total population was conserved at its ancestral size, dividing this at population splits to achieve equal populations in the most recent time period. All other parameters were unmodified. We ran 10 independent simulations using the same process described above with parameter draws allowing variation in divergence times and migration rates but not population sizes. Based on this data we calculated pairwise Fst and performed PCA using plink2. Results are shown in supplementary fig. S16.

### Signatures of selection based on extended haplotype homozygosity

We used haplotype-based methods to identify genomic regions under strong positive selection using our phased variant dataset. Specifically, we used the integrated haplotype score (iHS) to determine the regions under selection in individual populations by comparing the extended haplotype homozygosity (EHH) of major and minor alleles and used cross-population extended haplotype homozygosity (XP-EHH; XP-nSL) to compare the EHH patterns between a target population and the other populations. All test statistics were calculated using selscan v1.3.0 (Szpiech and Hernandez 2014) with default parameters. Next, the test statistics of iHS, XP-EHH, and XP-nSL were normalized in 50 separate allele frequency bins using the companion program norm. After normalization SNPs with extreme values were identified genome-wide based on the following criteria (|iHS|>2, XP-EHH/XP-nSL > upper first percentile). We then calculated the proportion of SNPs with extreme values within 50kb windows and identified windows as candidates for selective sweeps as those in the top 1% based on this proportion. This process was performed separately for each of the three test statistics (iHS, XP-EHH, XP-nSL) and multiIntersectBed (Quinlan and Hall 2010) was used to report the overlapping candidate regions of all tests.

Since our goal was to identify sweeps unique to each population we removed those that were significant based on iHS in more than one population. This was not required for the cross-population tests since those already target regions that differ between populations.

### Signatures of selection based on allele-frequency

Although haplotype-based methods (see above) formed our primary method for identifying selective sweeps, we also calculated population branch statistics (PBS) which measure the extent of differentiation in allele-frequencies between populations. First we used the --fst function in PLINK to calculate Fst statistics genome-wide for all pairs of populations, using the default Fst calculation (Hudson). These Fst values were then used to calculate the population branch statistic as described in (Yi et al. 2010).

### Calculation of empirical false discovery rate for signatures of selection based on population branch statistics

We used simulated data under our best-fitting demographic model with fastsimcoal2 to calculate the distribution of population branch statistics (PBS) for each population arising under neutrality. PBS values calculated on all simulated data include estimates from 50 simulations using randomly selected values across the bootstrap-estimated 90% confidence intervals for model parameters. Since this generated a much larger number of PBS values to our real dataset, and also includes many sites in LD we randomly selected 100k values from this simulated data and our real data. The resulting 200k were then ranked by PBS value (0 the highest) and the false positive rate for the ith ranked value was calculated by counting the number of false (ie simulated) values from ranks 0 through to i and dividing this value by 0.5i. We then calculated the threshold value, above which this empirically calculated error dropped below 0.01 (1%) and used this as our criteria for significance. This procedure was performed separately for each population.

### Mapping to pseudo-chromosomes

We used ragtag v.1.1.1(Alonge et al. 2019) to align the *Acropora digitifera* genome to the *Acropora millepora* chromosome-level genome assembly (Fuller et al. 2020) with default settings. This placed 735 of the 955 *A. digitifera* scaffolds in pseudo-chromosomes, comprising 97% of assembled bases. We used this mapping to translate between scaffold level and pseudo-chromosome coordinates for the purpose of visualization only. Specifically, it was used to create the Manhattan plot (fig 3A).

### Gene annotations

Gene models for the *Acropora digitifera* version 2 assembly were obtained from the authors of its original publication (Shinzato et al. 2020) in gene feature annotation (GFF3) format. As these gene models are based on scaffolds from the original assembly (available at https://marinegenomics.oist.jp/adig/viewer/info?project_id=87) that have not undergone the RefSeq curation process their coordinates needed to be updated to match the ncbi assembly (GCA_014634065.1) that we used for our analyses. To do this we first aligned the two genomes with Cactus (Armstrong et al. 2020) and then used the ucsc chain and liftOver utilities (Kuhn et al. 2013) to generate updated gene model coordinates. The resulting updated gene models and full details of the lift-over process are available via the online code repository https://github.com/bakeronit/acropora_digitifera_wgs for this paper.

Starting from these updated gene models we first selected the longest transcript per gene using cgat toolkit (Sims et al. 2014) and then extracted nucleotide and protein sequences for each coding sequence using gffread (Pertea and Pertea 2020). Functional annotations for these genes were then obtained by performing blastp and blastx searches on protein and nucleotide sequences respectively against the Swissprot database (downloaded 2021 May 9) (Bairoch and Apweiler 1996), filtering to include hits at e-value < 1e^−5^ only. We then selected the best available blast[xp] hit for each gene and assigned this as its closest putative homolog. In addition, we used the Uniprot ID mapping service to look up detailed functional information (including GO terms) for these homologs.

Our initial gene ontology enrichment analysis was performed based on these GO terms assigned based on blast hits to Swissprot, however, we found that this often resulted in enrichment of highly specific gene ontology terms that were clearly spurious as they involved functions that are not present in Cnidarians. To resolve this issue we decided to use GO terms assigned using Interproscan version 5.53-87 (Jones et al. 2014), which uses functional information assigned to conserved domains rather than to specific genes. A complete table of annotated genes resulting from both BLAST and Interproscan annotations is provided as supplementary table S9.

### GO enrichment analysis

Formal statistical analysis for enrichment of GO terms is challenging because the terms themselves are not independent, and because genes are not randomly distributed across the genome. The R package topGO v2.42 (Alexa et al. 2006) attempts to deal with the first of these issues (non-independence of GO terms) by weighting the assignment of genes to terms in a way that increases the significance of more specific terms at the expense of more biologically general parent terms. We, therefore, used topGO with the default “weight01” algorithm for all enrichment tests. To deal with the second issue (non random distribution of genes across the genome) we calculated enrichment statistics at two levels. First we evaluated enrichment at the gene level. In this analysis all genes overlapping with putative selective sweeps were assigned to the target set and the complete set of all annotated genes was assigned as the background set. Since this analysis ignores the fact that multiple genes from the same GO term might be present in the same sweep region we also performed an enrichment test based on sweeps rather than genes. As this test was used as a complement to the first we performed it only for GO terms that were significant at the gene level. To perform this second test we first assigned GO terms to all 50kb regions in the genome based on the GO terms assigned to overlapping genes. This analysis included both sweep regions and non sweep regions. A p-value based on Fisher’s exact test was then calculated by counting the number of sweep regions (a subset of all 50kb regions) with a given term and comparing this to the background count across all regions.

### Symbiont analysis

Although our sequencing strategy targeted the coral host, the resulting sequencing data contained a median of 260k reads classified by Kraken as originating from Symbiodiniaceae (minimum 4k, max 1.7M). We used several methods to quantify the symbiont diversity based on these reads. Firstly, a custom database composed of the genomes of five common coral associating Symbiodiniaceae genera and the *Acropora digitifera* genome assembly was built using kraken v1.0 (Wood and Salzberg 2014). Raw sequencing reads from all samples were then classified and assigned to each taxonomy. After confirming the dominance of *Cladocopium* in all samples, we tried to investigate the diversity within genera using three methods. Firstly, we mapped the reads to the mitochondrial genome of *Cladocopium* C1 and built a haplotype network using PopART (Leigh and Bryant 2015) with the consensus sequences of 41 samples after removing samples with less than 20X average mapping depth (excluding regions with no reads mapped) in Claocopium mitogenome. We also mapped non-host reads to ITS2 sequences from the symportal (Hume et al. 2019) database and quantified their abundance by counting the number of uniquely mapping reads to each ITS2 reference sequence. To make use of all symbiont reads regardless of genomic locus of origin, we performed an alignment-free method (https://github.com/chanlab-genomics/alignment-free-tools) to calculate the d2s metric based on shared k-mers in sequencing reads from each pair of samples. This produced a set of pairwise distances which we visualised using an MDS plot (fig 1E).

### Estimating the timing of selection at the peroxinectin locus

To investigate the timing of the selective sweep on the peroxinectin locus we used the R package starTMRCA (commit cf9f021 from github) (Smith et al. 2018) which estimates the time to the common ancestor (TMRCA) of haplotypes bearing a beneficial allele based on the length distribution of ancestral haplotypes and the accumulation of mutations since divergence. Since we did not know the beneficial allele, we instead identified alleles likely to be in complete linkage with the beneficial allele to serve as its proxy. We did this by choosing sites for which the derived allele was nearly fixed (on all but 3 haplotypes) in the inshore population and completely absent offshore. There were 84 such SNPs within the sweep locus, of which 75 were found within the gene s0150.g24 that overlapped with the strongest statistical indicators of selection (fig 4A). Of these 75 sites we chose 3 spanning the length of the gene (at positions 278594, 281245, 282923)

We then used VCFtools to export a 1Mb region centred on s0150.g24 from our phased vcf. For each of the 3 SNPs chosen as proxies for the beneficial allele we then used the R package REHH (Gautier and Vitalis 2012) to generate a furcation plot, and phytools (Revell 2012) combined with ggtree (Yu et al. 2018) to plot a midpoint rooted neighbour joining phylogenetic tree of the core haplotypes (central 200 sites). These visualisations all produced qualitatively similar results, all showing a clear distinction between selected and background haplotypes in the tree and strong extended haplotype homozygosity in the furcation plot.

We then ran starTMRCA separately for each of the 3 chosen SNPs using the 1Mb phased vcf as input. Other parameters were as follows; mutation rate of 1.2e^−8^ per base per generation, a recombination rate of 3.2e^−8^ per base per generation, chain length of 10000, proposal standard deviation of 20, initial value of TMRCA drawn from a uniform distribution from 0-10000 generations. Convergence was checked by running 10 independent chains and calculating the Gelman diagnostic using the coda package in R. For each SNP we recorded the median value of the posterior estimates of the TMRCA after discarding the first half as burn-in. Our final estimate for the time of selection on the locus is reported as the range of estimated values across these three SNPs.

The mutation rate used for starTMRCA analyses is the same as used for fastsimcoal2 and SMC++. The recombination rate was estimated based on a linkage map for *Acropora millepora* (Wang et al. 2009; Dixon et al. 2015) *which had a length of 1358 centimorgans. The rate used was then calculated by assuming a constant recombination rate and genome size of 430Mb for A. millepora*.

### Estimating allele age with GEVA

To estimate the time of origin for derived alleles in the peroxinectin locus we used Genealogical Estimation of Variant Age (GEVA) (Albers and McVean 2020). As GEVA requires polarisation of ancestral and derived alleles we performed this task first, using est-sfs (Keightley and Jackson 2018). Inputs to est-sfs were generated by performing a whole genome alignment of the A. digitifera genome to the genomes of two related species, Acropora millepora (GCF_013753865.1), and Acropora tenuis (http://aten.reefgenomics.org/) using progressive cactus v2.0.5 (Armstrong et al. 2020). We then updated our phased vcf to encode the ancestral allele as the reference allele and used this vcf as input to GEVA. GEVA was run assuming an effective population size of 10000, and used the same mutation rate used throughout (1.2e^−8^ per base per generation), and the same recombination rate (3.2e^−8^ per base per generation) as used for starTMRCA.

### Phylogenetic analyses of haem peroxidases

To investigate the evolutionary origins of the peroxinectin locus we used blastp to search for homologous genes in four other coral species, *Acropora millepora, Acropora tenuis, Porites lutea* and *Pachyseris speciosa*. Protein sequences for all genes identified as belonging to the haem peroxidase family (IPR019791) by Interproscan were extracted from *Acropora digitifera*. Using these as query sequences we identified all close homologs (e-value < 1e-10) from the protein sets of all other species using blastp. These were then aligned using the MAFFT (v7.394) (Katoh et al. 2002) with the algorithm set to auto. After masking positions with more than 50 missingness, IQ-Tree (v2.0.3; Nguyen et al. 2015) was used to perform tree inference based on this alignment with 1000 ultrafast bootstraps and automatic model selection using modelfinder.

## Acknowledgements

This work is supported by computational resources of the National Computational Infrastructure (NCI) National Facility systems through the NCI Merit Allocation Scheme (Project d85) awarded to C.X.C.

## Supplementary figures

Supplementary figures - AD

## Supplementary tables

Supplementary tables - AD

## Data availability

**Raw data:** NCBI PRJNA805369

**Code and accessory data:** https://github.com/bakeronit/acropora_digitifera_wgs

## Notes

https://github.com/bakeronit/acropora_digitifera_wgs

### Competing Interest Statement

The authors have declared no competing interest.

