## Supplementary Figures for "Evolutionary responses of a reef-building coral to climate change at the end of the last glacial maximum"

### Supplementary Materials

### Founder effects and adaptive selection drive rapid post-glacial divergence in a reef-building coral

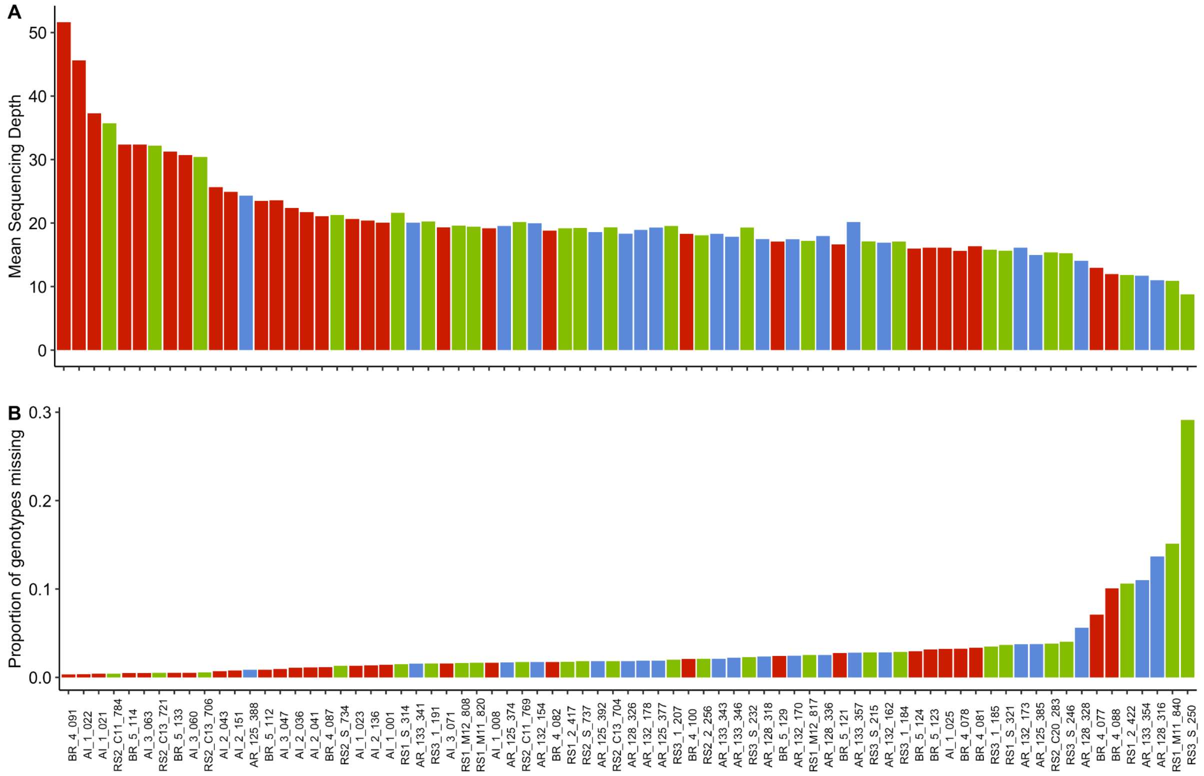


**Supplementary figure S1:** Mapping depth (A) and genotype missingness (B) of all samples. The colour of dots represents the sample origin population, inshore (red), north offshore (blue), and south offshore (green).

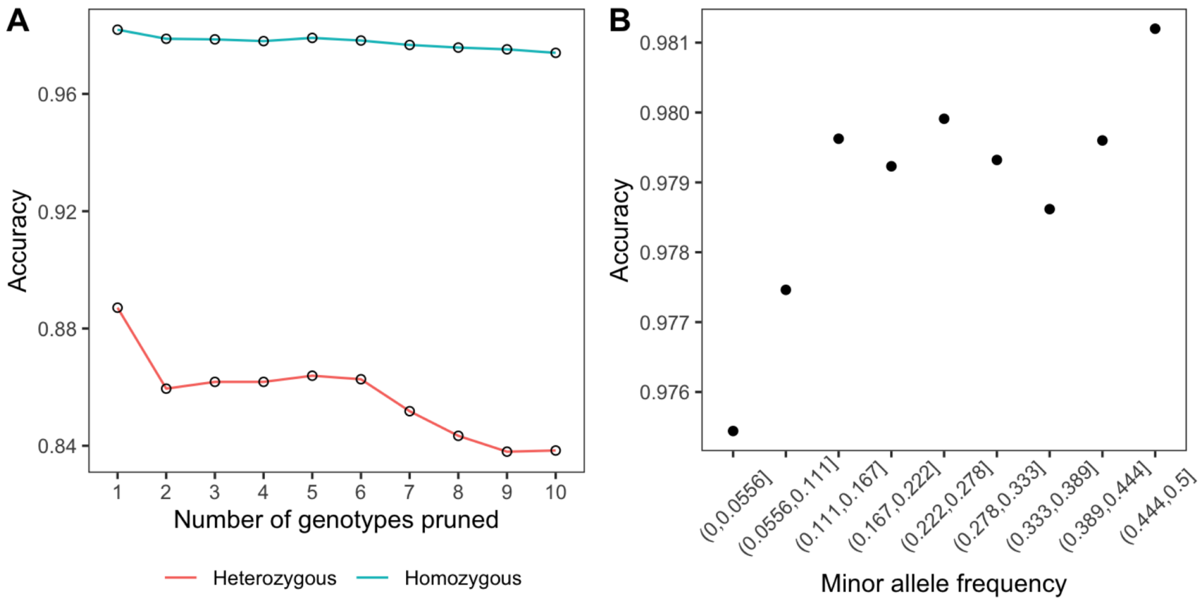


**Supplementary figure S2:** A. The estimated imputation accuracy at homozygous and heterozygous sites as a function of the number of missing genotypes. B. Estimated imputation accuracy as a function of minor allele frequency.

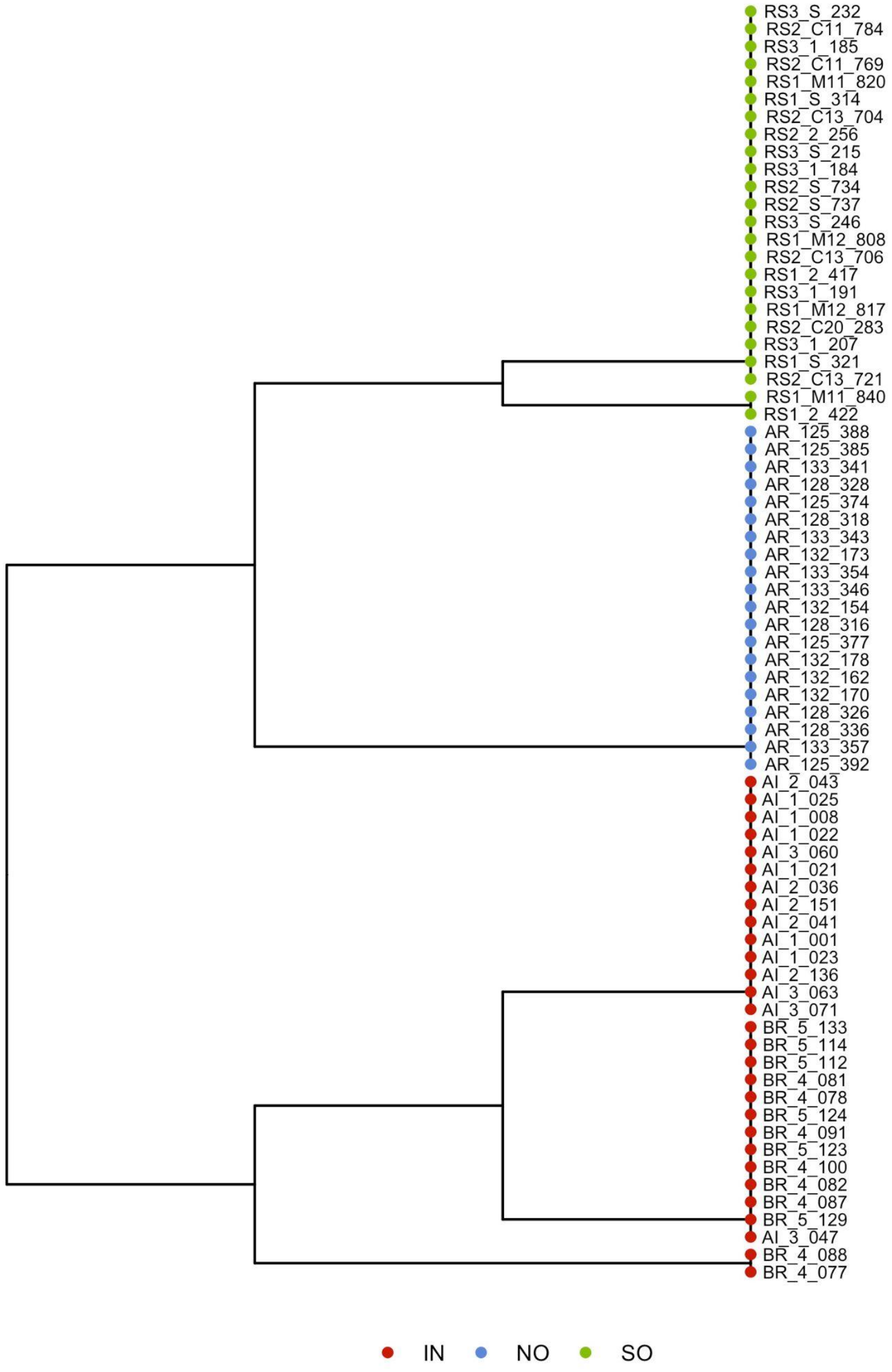


**Supplementary figure S3:** Sample tree inferred by fineStructure showing nodes with greater than 99% bootstrap support. Samples are coloured according to the location using the standard scheme (fig 1 in the main text).

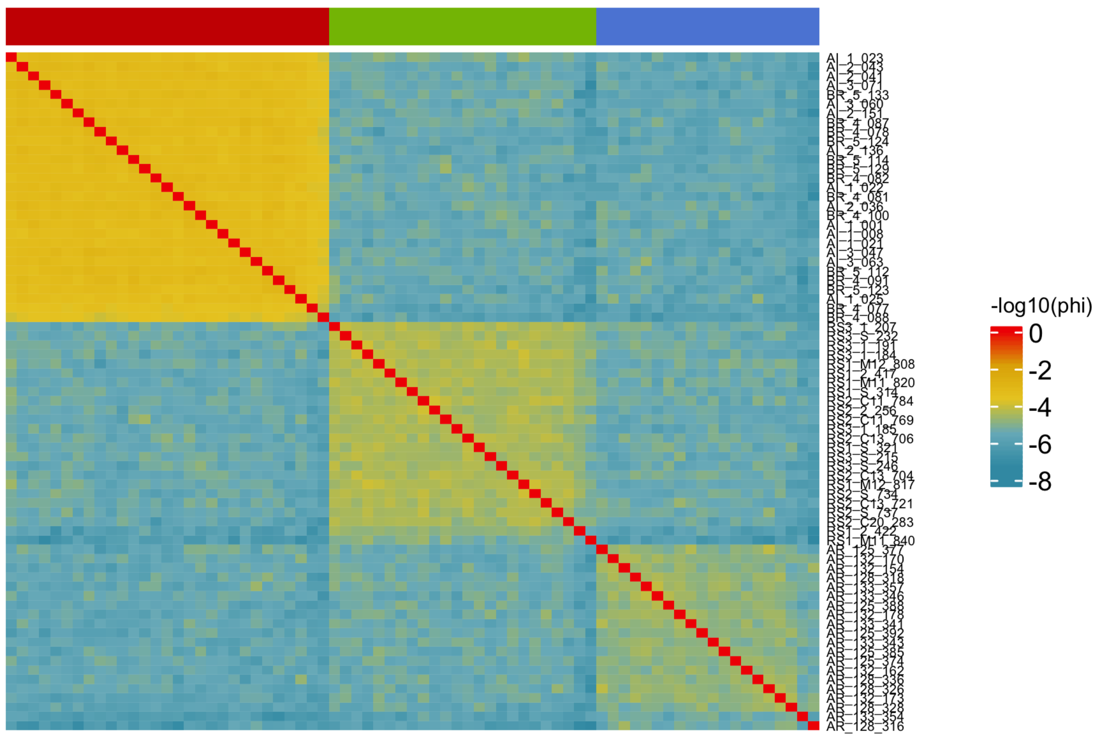


**Supplementary figure S4:** Clustered heatmap based on pairwise relatedness (phi) inferred by the relative proportion of IBD (identical by descent) segments.  Rows and columns both represent samples and use the same clustering. Sample labels are shown for rows and column colours indicate their location using our standard colour scheme (fig 1 main text).

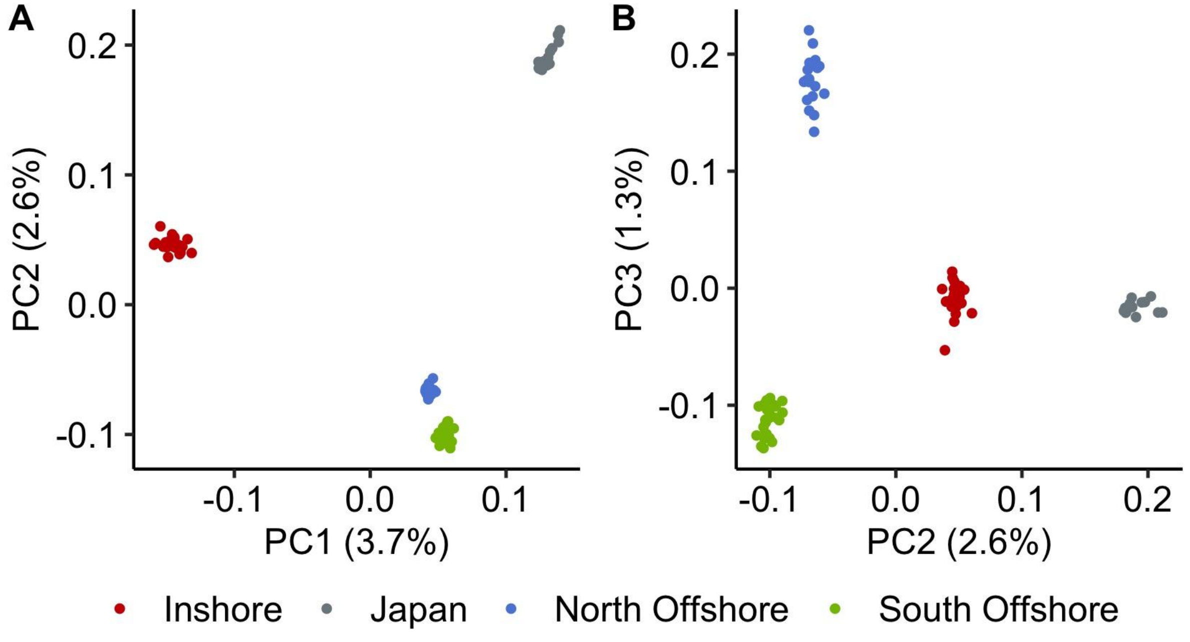


**Supplementary figure S5:** PCA of *A. digitifera* from Kimberley, Western Australia and Ryukyu Archipelago Japan, based on allele frequency. Plot (A) shows PC1

against PC2 and (B) shows PC2 against PC3. Samples are coloured according to location using our standard colour scheme (fig1 main text) with the addition of grey to indicate samples from Japan.

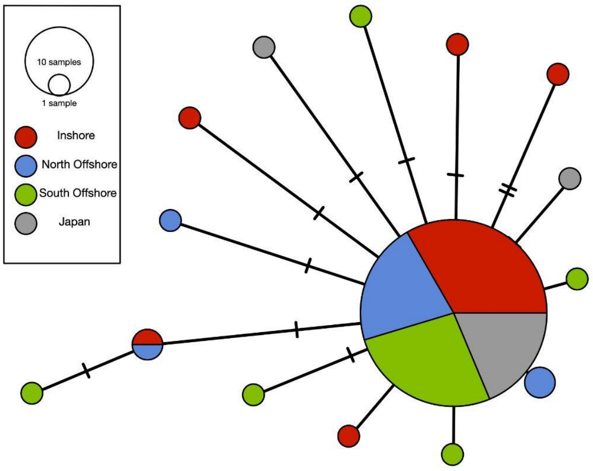


**Supplementary figure S6:** Haplotype network based on mitochondrial genomes of *A. digitifera* samples from the Kimberley region, Western Australia, and the Ryukyu Archipelago, Japan. Cross bars on edges indicate the number of mutations separating haplotypes while the size of nodes indicates the number of samples with the same haplotype.

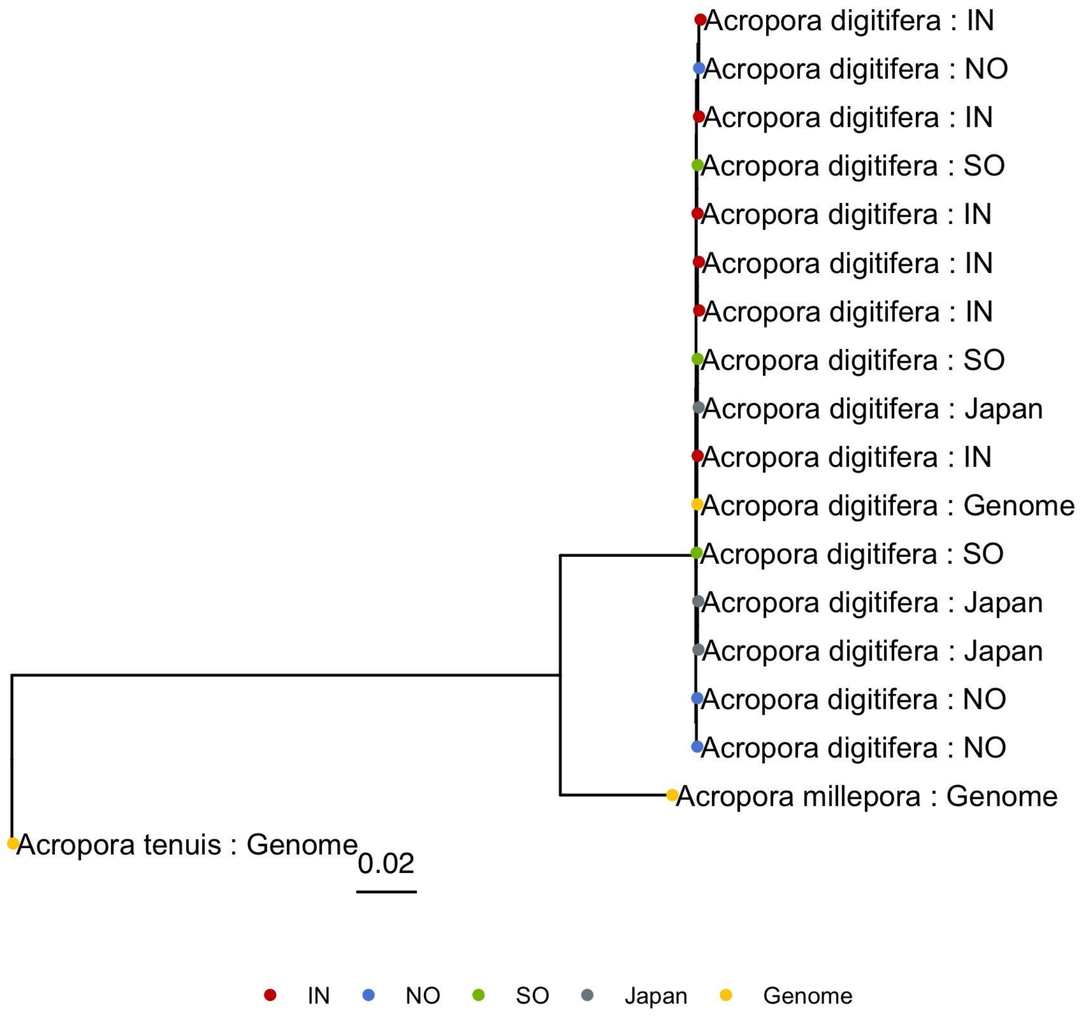


**Supplementary figure S7:** Maximum likelihood tree inferred from consensus UCE and Exon sequences. Sequences include those obtained from published reference genomes for *Acropora millepora* (GCF_004143615.1; (Ying et al. 2019)), *Acropora digitifera* (GCA_014634065.1, (Shinzato et al. 2020)), and *Acropora tenuis* (http://reefgenomics.org/aten/; (Cooke et al. 2020)) as well as representative population genomic samples from our study (NO, SO, IN) and from Japan (NCBI Bioproject PRJDB4188; (Shinzato et al. 2015)). Japanese samples correspond to SRA accessions (DRR099286, DRR099287, DRR099291). Samples from our study included (IN: AI_1_001, AI_1_008, AI_1_021, BR_4_077, BR_4_078, BR_4_081; NO: AR_125_374, AR_125_377, AR_125_385; SO: RS1_2_417, RS1_2_422, RS1_M11_820).

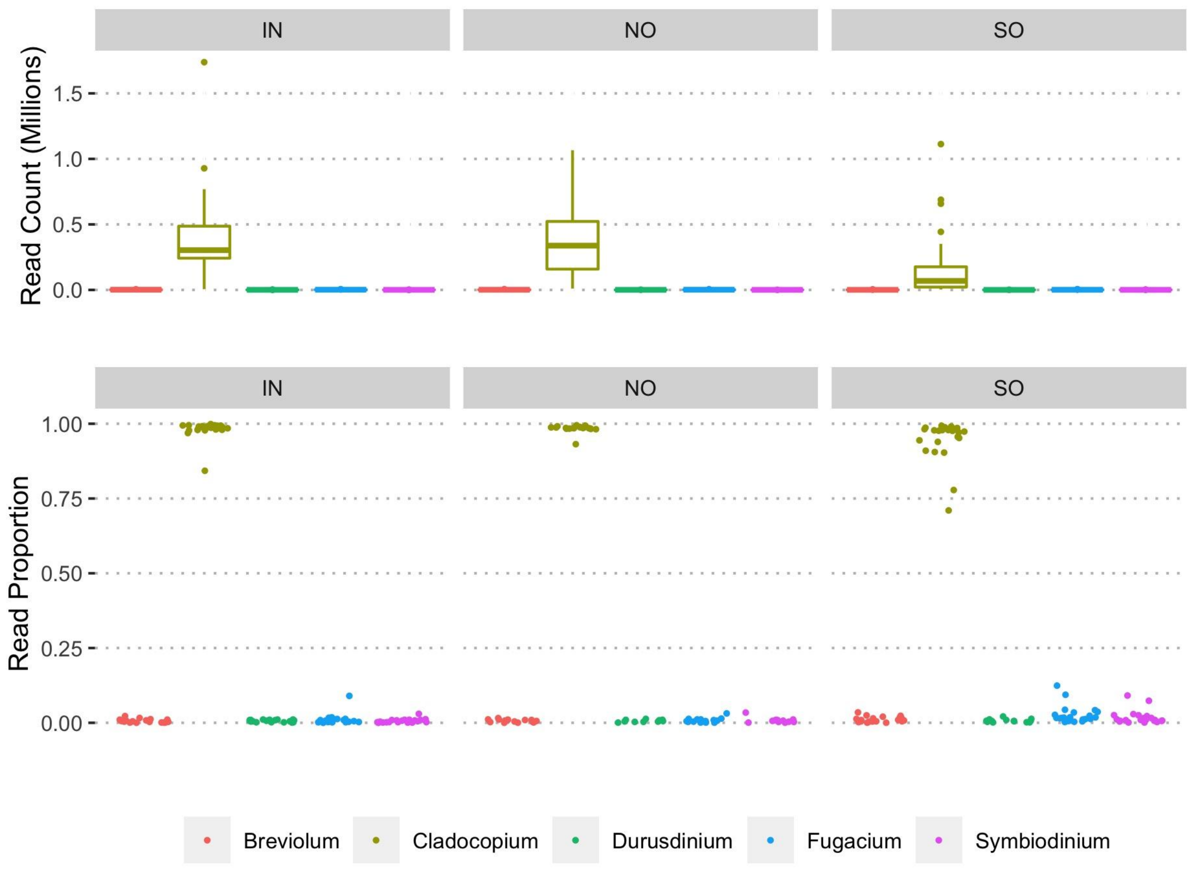


**Supplementary figure S8:** Summary of reads classified as Symbiodiniaceae using Kraken. Both top and bottom plots show the spread of values measured across individual samples. Read counts are shown as sample totals (top) and as proportion of the sample total (bottom) across five genera of Symbiodiniaceae. Samples are plotted separately for each location (IN: Inshore, NO: North Offshore, SO: South Offshore).

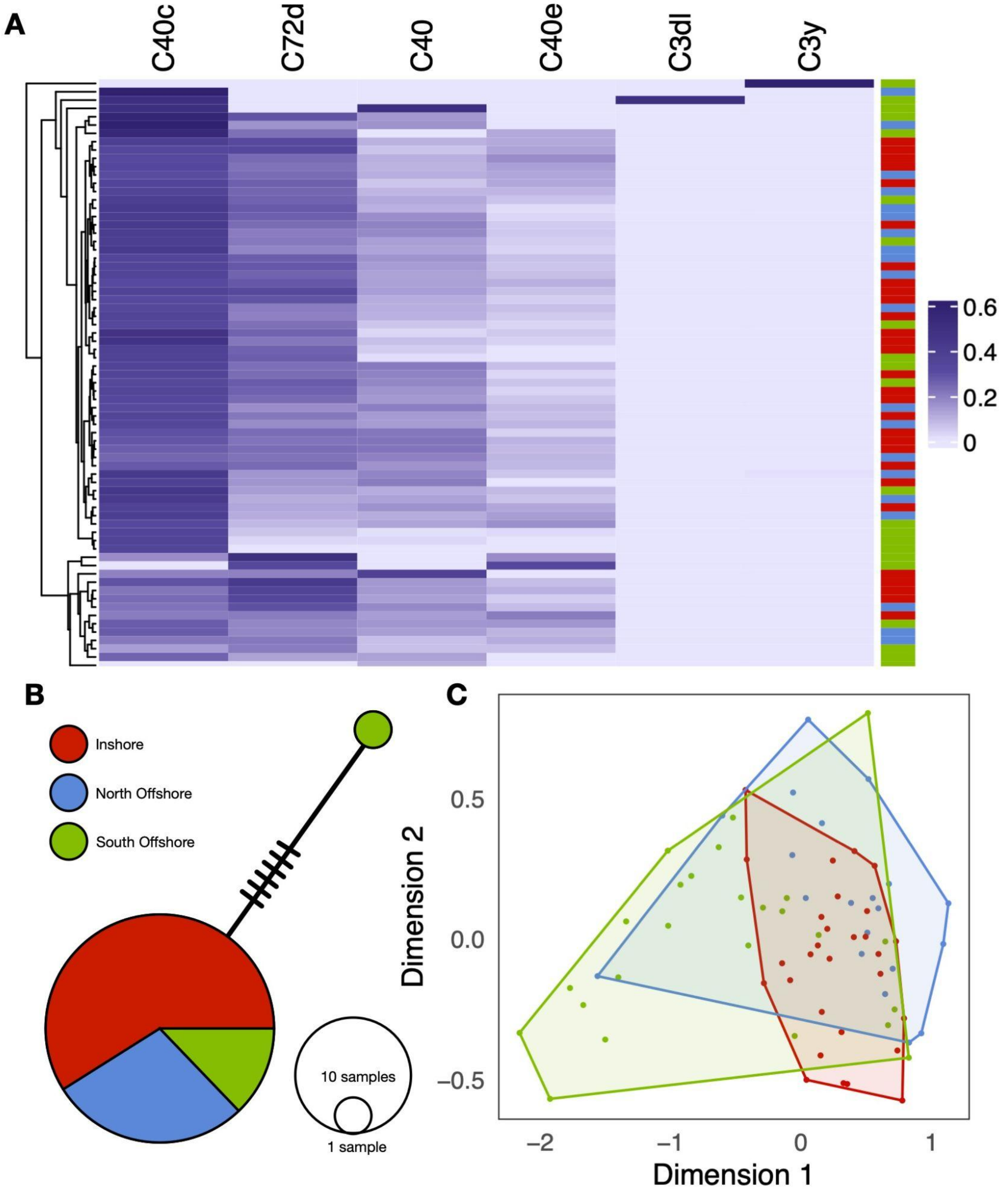


**Supplementary figure S9:** Diversity of sequences related to the dominant symbiont genus, *Cladocopium*. A. Heatmap of read counts mapping to symportal ITS2 reference type sequences. Rows represent coral samples, and the columns show the detected ITS2 types from read mapping. Coloured strip on the right indicates the location of origin for each sample using the colour scheme shown in B. B. Haplotype network based on mitochondrial sequences for 41 samples for which sufficient reads were available to allow consensus calling.  Edge cross bars indicate the number of mutations separating haplotypes and the size of nodes indicates the number of samples with a given haplotype. C. Multidimensional scaling (MDS) plot based on pairwise distances between samples calculated using D2S statistics. D2S statistics are calculated based on kmer counts in reads of *Cladocopium* origin (that map to the *Cladocopium goreaui* genome). Convex hulls enclose points from each location and are coloured according to our standard location color scheme (see B).

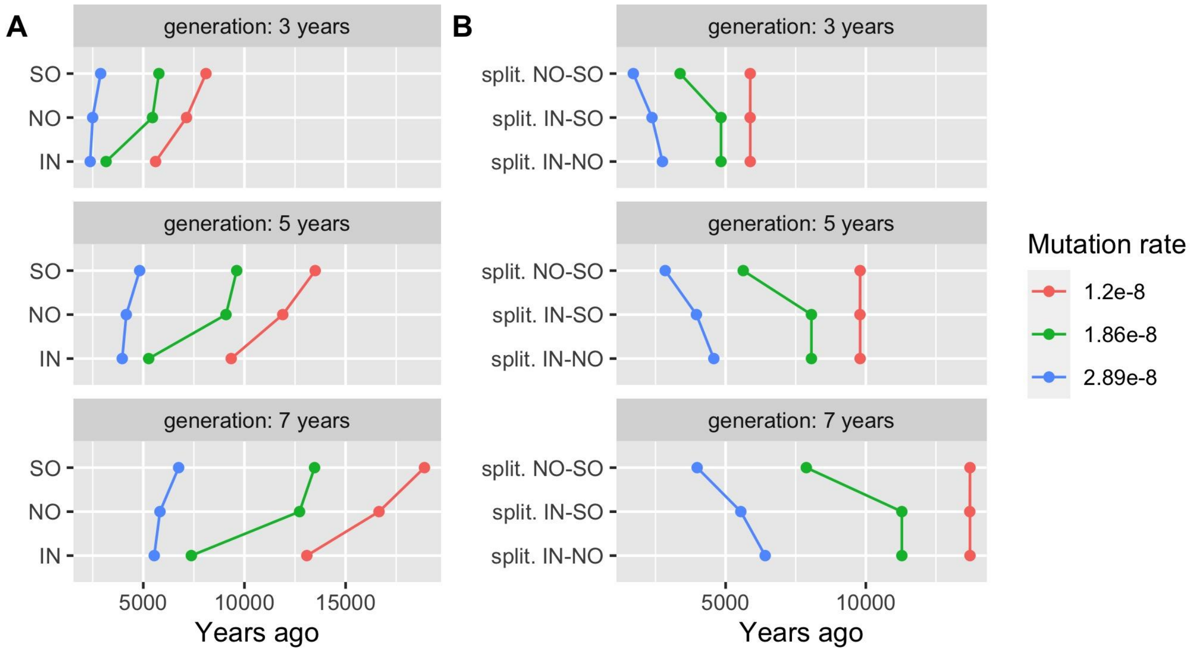


**Supplementary figure S10**: Variation in the estimated timing of key demographic events under different mutation rates and generation times. All estimates were obtained using SMC++. A. the bottleneck time of inshore (IN), north offshore (NO), and south offshore (SO) *A. digitifera* populations. B. the split time between each pair of populations.

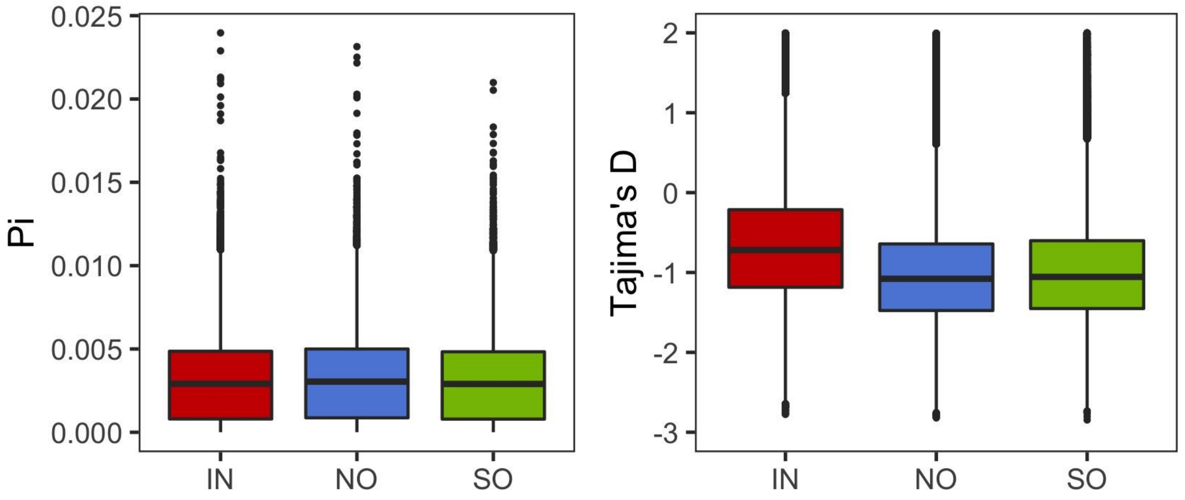

**Supplementary figure S11:** Boxplots showing the genome-wide distribution of nucleotide diversity (Pi; left) and of Tajima's D (right). Both plots show results for each of the three populations separately and use our standard color scheme to denote location (fig 1 main text).

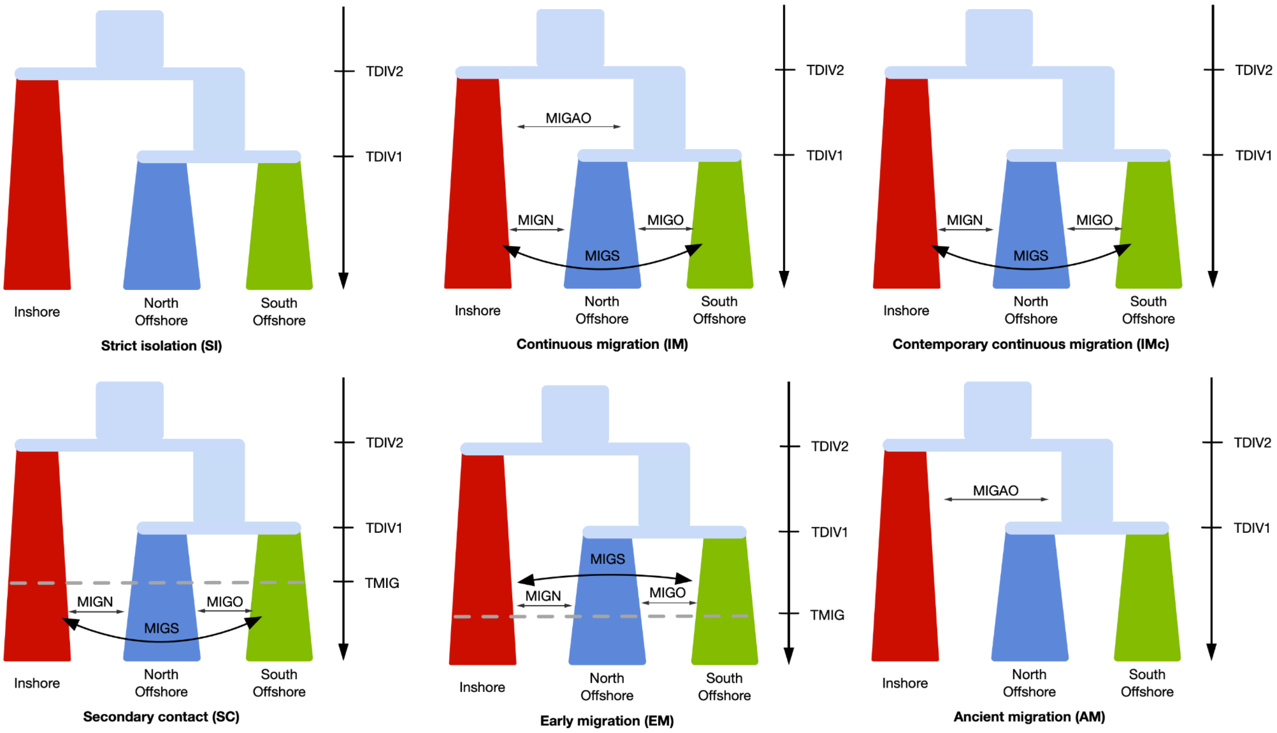


**Supplementary figure S12:** Schematic diagram of six alternative demographic models used in fastsimcoal2. Time is shown from most ancient (top) to present day (bottom). All models were allowed to have an exponential growth rate. The parameters TDIV1 and TDIV2 represent the time of offshore-offshore divergence and inshore-offshore divergence, respectively. Moving forward in time, TMIG represents the time at which migration starts. In models with TMIG there is no migration prior to TMIG. In models without TMIG the migration parameters persist only during one of the time intervals delineated by TDIV1 and TDIV2.

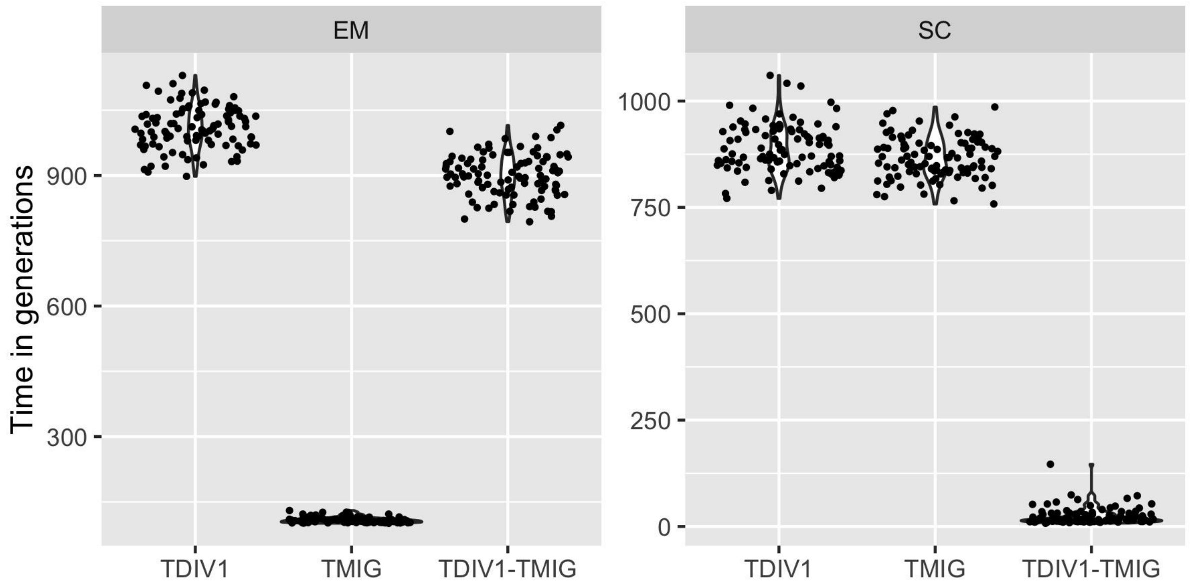


**Supplementary figure S13:** Estimates from fastsimcoal2 for the time of offshore-offshore divergence (TDIV1) and the migration start time (TMIG) in models EM and SC (see figure S12). Points are randomly jittered on the x-axis to avoid overplotting and show the distribution over 100 independent fastsimcoal2 runs.

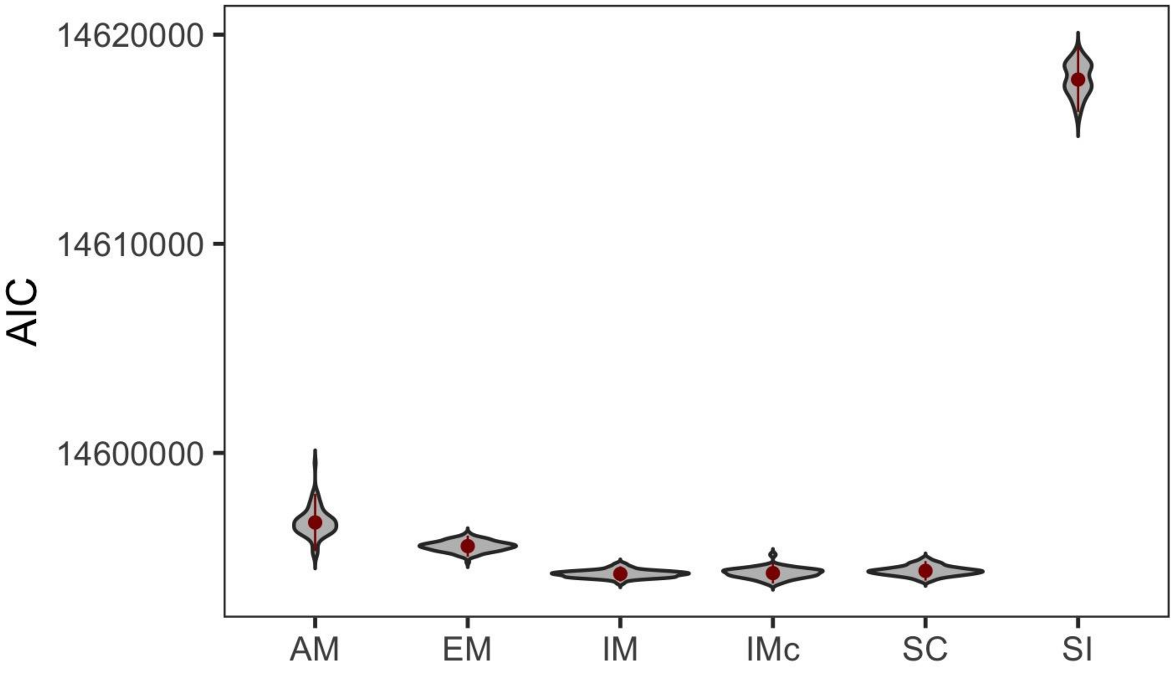


**Supplementary figure S14:** Distributions of AIC values from 100 independent fastsimcoal2 runs for each model described in supplementary figure S12.

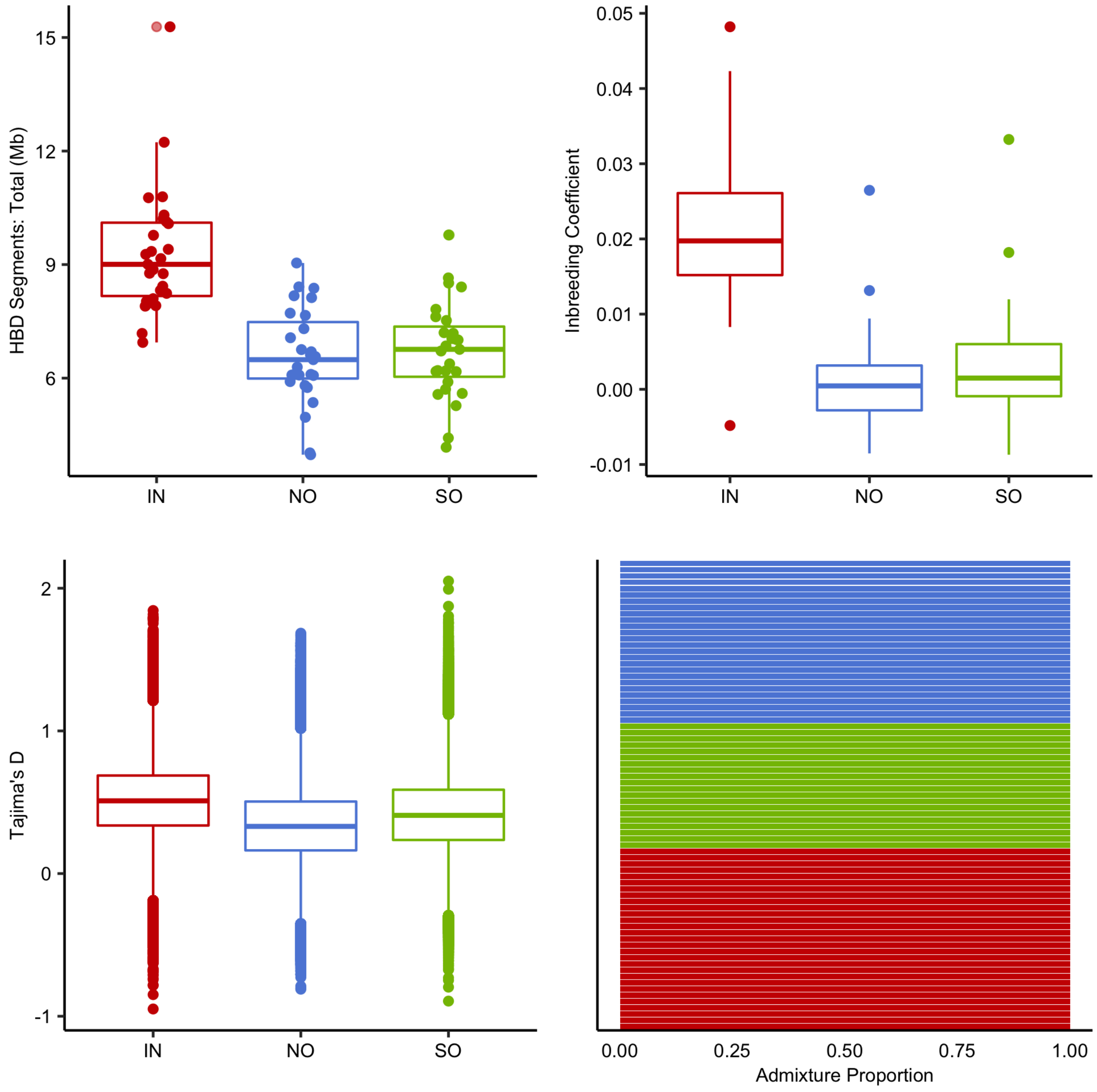


**Supplementary figure S15:** Population genetic statistics and Admixture proportions calculated based on simulated data under the best fitting model from fastsimcoal2 (model IMc in supplementary table S8). Sample locations are named and coloured according to our standard scheme (main text fig 1). Boxplots show the distribution of values from 50 simulation runs with fastsimcoal2 based on independent draws across the error range of model parameters. A single representative Admixture plot is shown as all simulations produced near-identical plots.

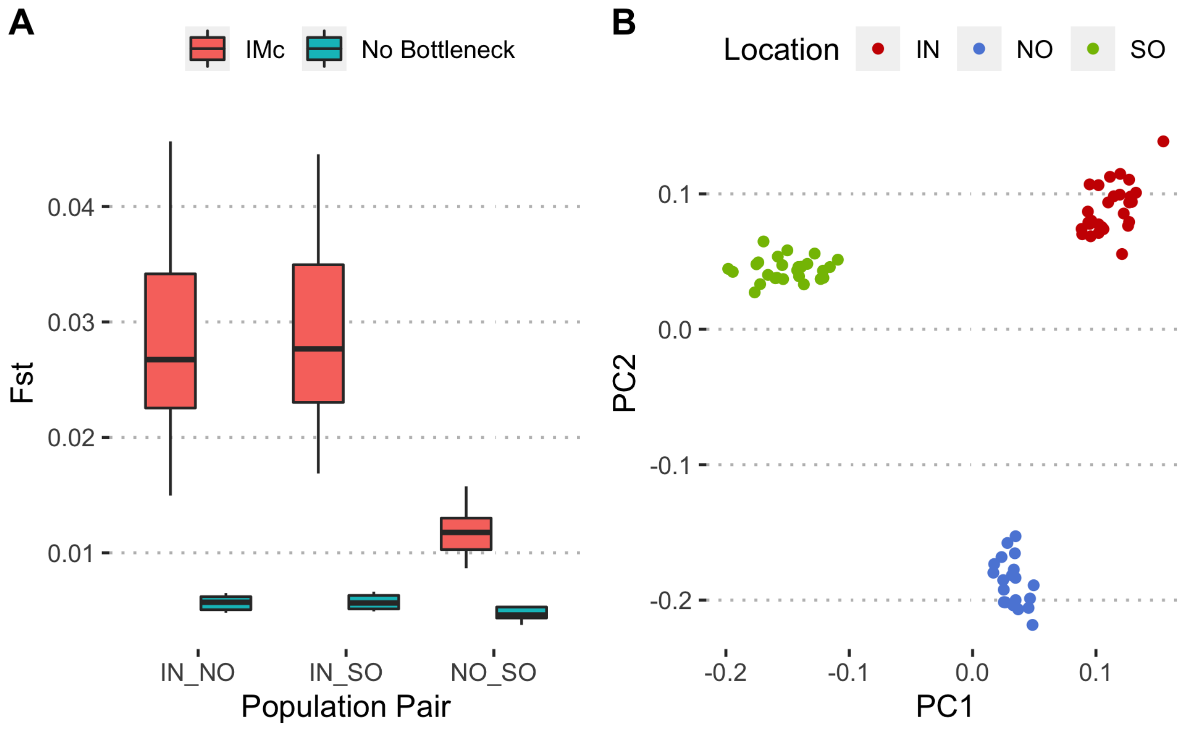


**Supplementary figure S16:** Influence of the bottleneck on population structure and divergence. **A.** Boxplots of pairwise Fst (Hudson) for the full IMc model compared with a model with constant population size (No Bottleneck). Spread of values is from 10 independent simulations for each model. **B.** PCA showing population structure under the No Bottleneck model. PCA shows results for one simulation. Other replicates displayed qualitatively similar results.

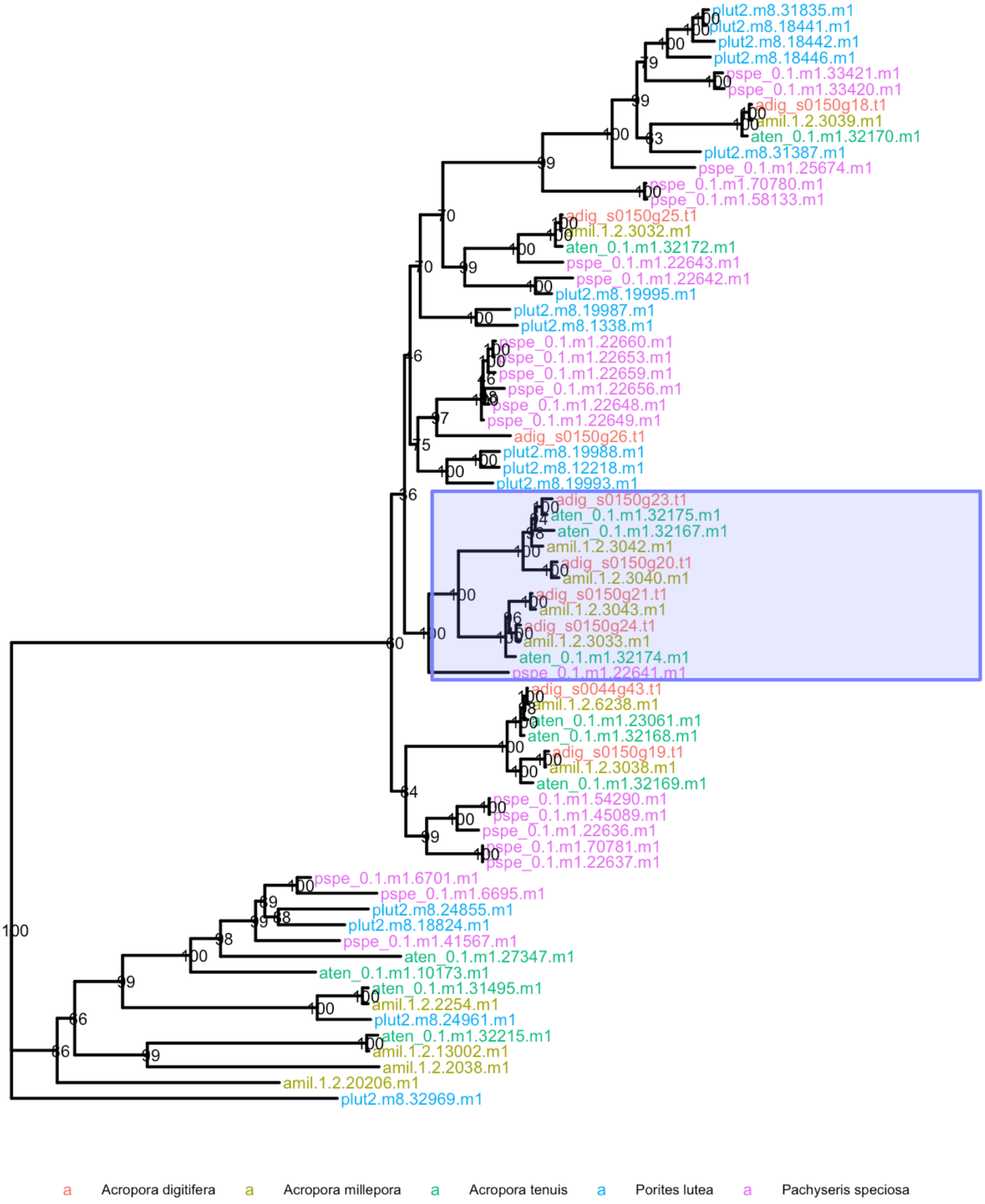


**Supplementary figure S17:** Phylogenetic relationships among haem peroxidases in representative coral genomes. Species chosen include three representatives of the genus Acropora and two outgroups, *Porites lutea* and *Pachyseris speciosa*. Highlighted clade includes four genes from the peroxinectin locus in A. digitifera that was examined in detail in the main text. All genes within the highlighted clade form clusters (closely spaced within the genome) in their respective species. The phylogeny shown is a subtree of the full phylogeny of haem peroxidases that includes all 8 members of the co-located peroxinectin cluster in *A. digitifera* as well as an outgroup used to root the tree. Nodes show bootstrap values based on 1000 ultrafast bootstrap replicates in IQ-Tree.

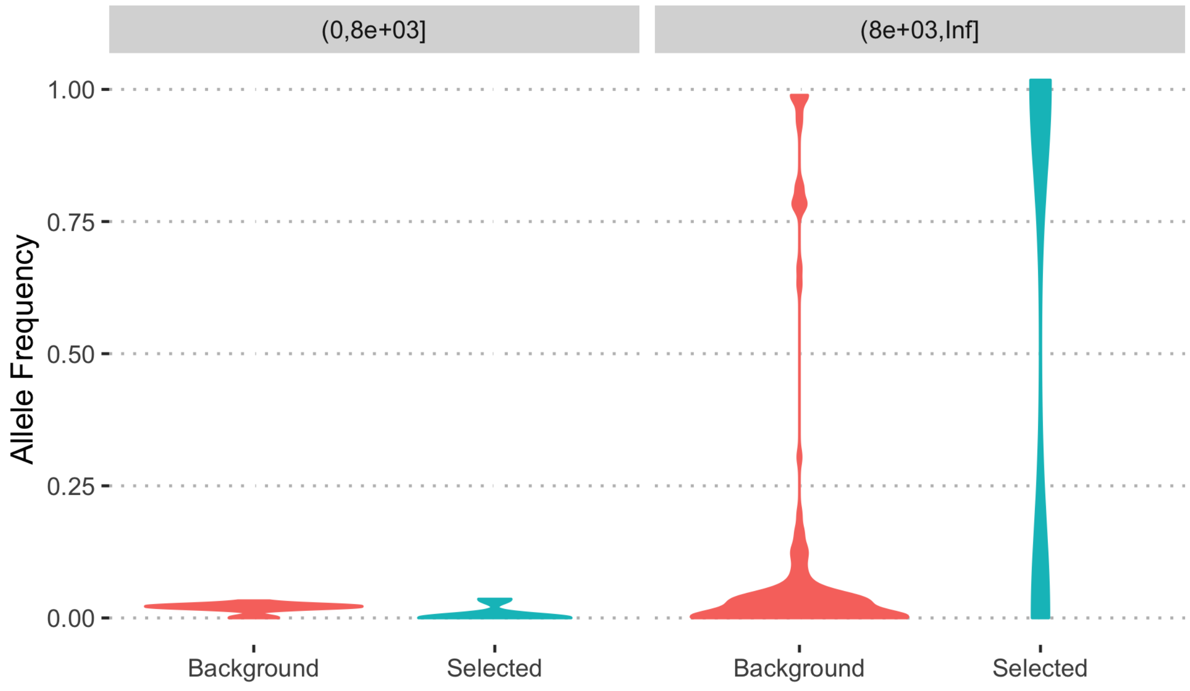


**Supplementary figure S18:** Relationship between frequency and age for derived alleles at 157 SNPs within the gene s0150.g24. Distributions of allele frequencies are shown using violin plots and split by age class (0-8Kya : left) and (>8Kya : right). For each allele its frequency in background haplotypes (red) is calculated separately from in selected haplotypes (blue).
